# An iPSC-derived small intestine-on-chip with self-organizing epithelial, mesenchymal and neural cells

**DOI:** 10.1101/2024.01.04.574203

**Authors:** Renée Moerkens, Joram Mooiweer, Aarón D. Ramírez-Sánchez, Roy Oelen, Lude Franke, Cisca Wijmenga, Robert J. Barrett, Iris H. Jonkers, Sebo Withoff

**Affiliations:** Department of Genetics, University of Groningen, University Medical Center Groningen, Groningen, The Netherlands; Oncode Institute, Utrecht, The Netherlands; Board of Governors Regenerative Medicine Institute, Cedars-Sinai Medical Center, Los Angeles, CA, USA; F. Widjaja Foundation Inflammatory Bowel Disease Institute, Cedars-Sinai Medical Center, Los Angeles, CA, USA

**Keywords:** induced pluripotent stem cell, organ-on-chip, intestine-on-chip, gut-on-chip, small intestine, human, intestinal epithelial barrier, mesenchyme, enteric neuron, differentiation medium

## Abstract

Human induced pluripotent stem cell (hiPSC)-derived intestinal organoids are valuable tools for researching developmental biology and personalized therapies, but their closed topology and relative immature state limits their applications. Here we use organ-on-chip technology to develop a hiPSC-derived intestinal barrier with apical and basolateral access in a more physiological in vitro microenvironment. To replicate growth factor gradients along the crypt– villus axis, we locally exposed the cells to expansion and differentiation media. In these conditions, intestinal epithelial cells self-organize into villus-like folds with physiological barrier integrity and myofibroblast and neural subtypes emerge and form a layer in the bottom channel underneath the epithelial tissue. The growth factor gradients efficiently balance dividing and mature cell types and induce an intestinal epithelial composition, including absorptive and secretory lineages, resembling the composition of the human adult small intestine. The result is a well-characterized hiPSC-derived intestine-on-chip system that can facilitate personalized studies on physiological processes and therapy development in the human small intestine.

## Introduction

The small intestinal epithelial barrier plays a pivotal role in sustaining intestinal homeostasis by preventing commensal microbes and harmful agents from entering the lamina propria while allowing absorption of nutrients and water. This intricate function is facilitated by the diverse intestinal epithelial cell types that form the barrier and which are constantly renewed. The local microenvironment, containing growth factors provided by neighboring mesenchymal cells, is essential to guide the continuous epithelial cell proliferation and differentiation processes along the crypt–villus axis^1^. Human-based model systems that capture the diversity of the small intestinal barrier are highly relevant for modeling processes like nutrition and drug metabolism and uptake, interactions with the microbiome and mucosal immune cells, and diseases such as celiac disease and inflammatory bowel disease.

Intestinal organoids successfully emulate many aspects of the human small intestinal epithelial barrier, but the closed topology of organoids complicates studies into barrier function and complex co-cultures of multiple cell types^2^. Moreover, preserving both the proliferative and mature cell types in human intestinal organoids has been challenging because this requires local activation and inhibition of several developmental pathways (e.g. WNT, BMP, EGF and Notch) in the form of growth factor gradients^3^.

Microfluidic intestine-on-chip systems have the potential to overcome these limitations as they contain multiple compartments that provide access to the apical and basolateral side of the epithelial barrier, allowing exposure to different microenvironments on either side of the tissue^4^. This compartmentalization also provides more control in the introduction of multiple cell types (e.g. microbial or immune cells) and enables the study of cell–cell interactions^5^. The continuous fluid flow through the microchannels of intestine-on-chip systems imposes physiologically relevant strain on the cells, which mimics luminal shear stress and promotes self-organization of the tissue^6,7^.

Most studies on steering human intestinal epithelial cells toward a specific lineage or physiological composition use adult stem cell-derived tissues, either as organoids or in static or microfluidic scaffold-based systems^1,8–10^. These studies have yielded widely used compositions of expansion media and several types of differentiation media, depending on the desired lineage. In contrast, only a few studies have used differentiation-type media to direct epithelial subtype development in human embryonic or induced pluripotent stem cell (hiPSC)-derived intestinal tissues^11^.

hiPSC-derived intestinal tissue can be generated from cells obtained by minimally invasive means (e.g. blood, urine or skin samples), thereby providing wide availability of case and control biomaterial, for example from large biobanks^12^. Additionally, as they are pluripotent, hiPSCs provide an unlimited resource for generating multiple tissues, which allows the study of organ–organ or organ–immune interactions in the same genetic background. The differentiation of hiPSCs toward intestinal epithelial organoids includes the co-development of mesenchymal cells, which increases the complexity of these tissues^13^. Previous efforts to increase the maturation and cell-type diversity of hiPSC-derived intestinal tissues mainly included transplantation into mice^11,14–16^, which limits applications. However, the application of a physiological microenvironment that includes relevant shear stresses^17^ or growth factor gradients *in vitro* is largely unexplored.

We aimed to create a model system of the human small intestinal barrier by exploiting the advantages of hiPSC-derived tissues, intestine-on-chip systems, and dual expansion and differentiation media exposure to replicate intestinal growth factor gradients and balance proliferating and mature cell types. Within our intestine-on-chip, we identify epithelial, mesenchymal and neural populations that self-organize into villus-like folds and subepithelial tissue, respectively. By combining single-cell RNA-sequencing (scRNAseq), microscopy and flow cytometry, we provide a high-resolution characterization of cell-type composition and demonstrate a close resemblance to the cell types and composition in the human small intestine.

## Results

### Development of a hiPSC-derived intestine-on-chip with physiological growth factor gradients

We aimed to develop a hiPSC-derived intestine-on-chip with physiologically relevant small intestinal epithelial cell-type composition by providing culture media that resemble the conditions along the crypt–villus axis in the human small intestine. Intestinal epithelial organoids were differentiated from three hiPSC lines based on previously published protocols^17,18^. EPCAM-positive intestinal epithelial cells were enriched using immunomagnetic positive selection to remove the majority of mesenchymal cells that emerged during the differentiation. After selection, the cell suspension contained 97% (standard deviation (SD) 2.2) EPCAM-positive intestinal epithelial cells and 3% (SD 2.0) EPCAM-negative cells. This cell suspension was seeded into the top channel of the Emulate Chip-S1, a microfluidic system containing two channels separated by a porous membrane^19^, and grown for 7 days in an expansion medium (EM) under a constant flow over the top and bottom channel to establish a tight barrier (Days −7 to 0; Figure 1A). During this time, the monolayer of intestinal epithelial cells on the membrane self-organized into three-dimensional villus-like folds and the permeability to 3-5 kDa fluorescein isothiocyanate– dextran (FITC-dextran) stabilized at a value (0.11 x10^-6^ cm/s (SD 0.03 x10^-6^)) remarkably similar to that of the human small intestine (∼0.11-0.12×10^-6^ cm/s) (Figures 1B, 1C and S1A)^20^. The villus-like folds only formed in the top channel where top and bottom channels were aligned (Figure 1D).

**Figure 1.**
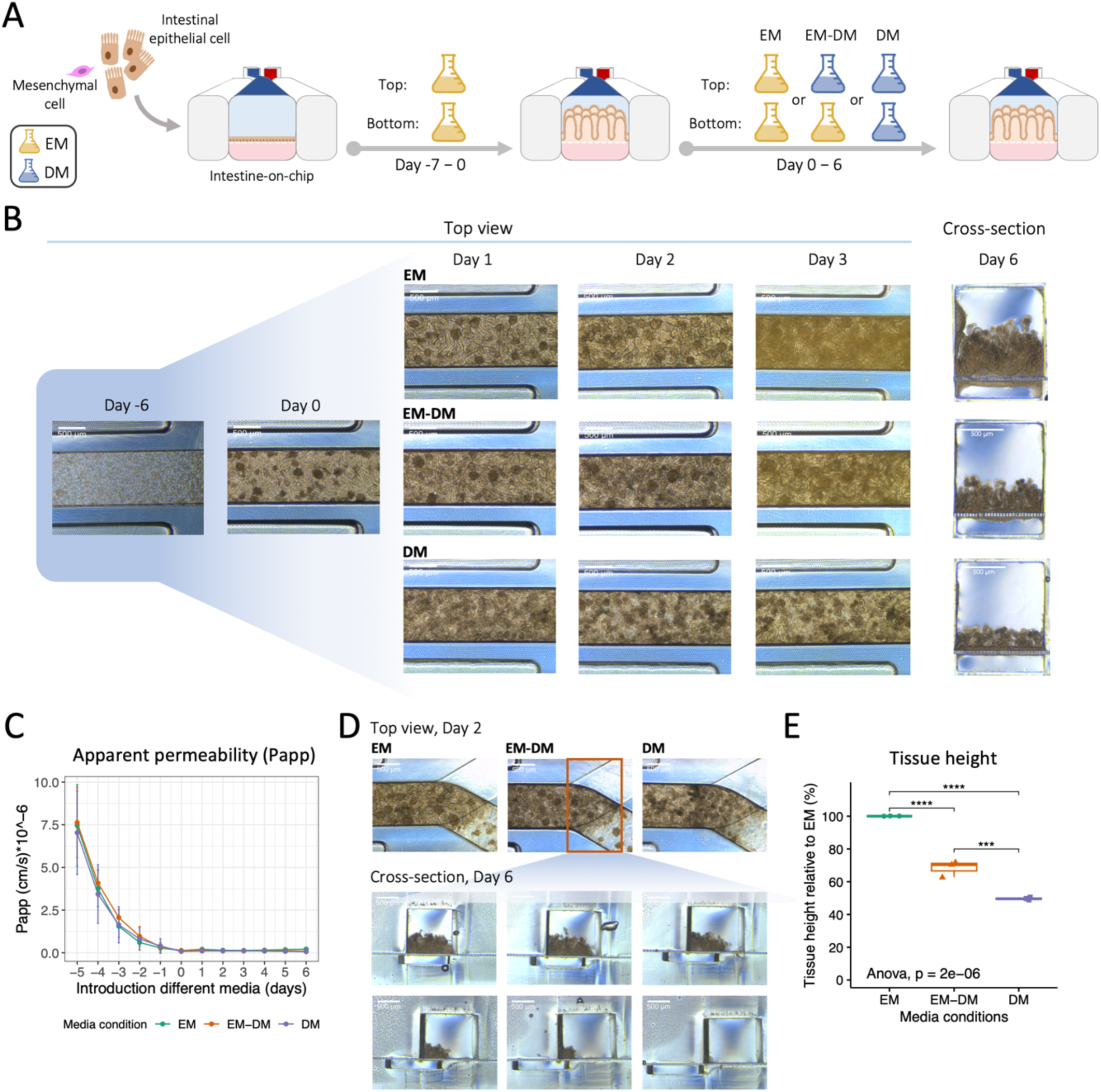
Media conditions influence tissue morphology in the intestine-on-chip. (A) Schematic overview of experimental design and timeline. EM = expansion medium, DM = differentiation medium. (B) Representative brightfield images of the top and cross-sectional view of the intestine-on-chip over time. (C) Apparent permeability of the tissue to 3-5 kDa fluorescein isothiocyanate–dextran, with the top channel as the dosing channel and the bottom channel as the receiving channel, displayed as average values from three biological replicates with standard deviation. (D) Representative brightfield images of the top and cross-sectional view of the intestine-on-chip on the intersection of the top and bottom channel. (E) Quantification of tissue height in the top channel in different media conditions, expressed relative to tissue height in the EM condition. Median values from three biological replicates are shown. P-value ≤ 0.05 (*), 0.01 (**), 0.001 (***), 0.0001 (****).

Next, to simulate the growth factor gradients along the crypt–villus axis in the human small intestine, we introduced a differentiation medium (DM) lacking WNT activators and BMP inhibitors and containing Notch- and MAPK pathway inhibitors to the top channel, while continuing exposure to the proliferation-inducing EM in the bottom channel. The intestine-on-chip exposed to this dual condition (EM-DM) was compared to systems with only EM or DM in both channels (Days 0 to 6; Figure 1A). Already at day 1, we could observe condition-specific differences in villus-like fold density, and these gradually increased over time (Figures 1B and S2). At day 6, the end of the experiment, the tissue height and cell density were highest in EM (522 μm (SD 125) = 100%, 3.7 x10^6^ cells (SD 1.0 x10^6^)), substantially lower in EM-DM (68.4% (SD 5), 1.4 x10^6^ cells (SD 0.3 x10^6^)) and lowest in DM (49.7% (SD 0.8), 0.9 x10^6^ cells (SD 0.3 x10^6^)) (Figures 1E, S1B and S1C). Regardless of the differences in morphology and cell density, the permeability to FITC-dextran remained stable from days 0 to 6 in all conditions (0.13 x10^-6^ cm/s (SD 0.04 x10^-6^)) and the viability of cells after dissociation at day 6 (91% (SD 0.5)) was not affected by the media conditions (Figures 1C and S1D). The morphological differences we observed were consistent with continuous proliferation in EM, induction of differentiation in DM and balanced proliferation and differentiation in the EM-DM dual condition.

### Epithelial and mesenchymal cells self-organize and differentiate in the intestine-on-chip

We observed that, even though cells were only seeded in the top channel of the intestine-on-chip system, they also emerged in the bottom channel during days −7–0 and expanded regardless of media condition during days 0–6 (Figures 1B, S1A and S2). To characterize the cell-type composition in both channels in the different media conditions, we performed immunofluorescent staining on cross-sectional slices and flow cytometry on dissociated cells using intestine-on-chip systems cultured until day 6. In all media conditions, the cells in the top channel were predominantly intestinal epithelial cells (EPCAM-and CDX2-positive), whereas the cells in the bottom channel were VIM-positive and EPCAM*-* and CDX2-negative cells, suggesting that these cells represent mesenchymal cells (Figures 2A, S3A and S3B)^21^. Mesenchymal cells that were localized in the pores of the membrane, separating the top and bottom channel, were detected in all media conditions, demonstrating their capacity to migrate from the top to the bottom channel (Figure S3C). The localization of mesenchymal cells relative to intestinal epithelial cells may reflect their natural position in the human small intestine, where (myo)fibroblasts form a supportive layer directly under the intestinal epithelial barrier^22,23^. Mesenchymal cells were most prominent in the EM condition, whereas they were reduced and restricted to the bottom channel in the EM-DM condition and almost depleted in the DM condition (Figures 2A, S3A and S3D).

**Figure 2.**
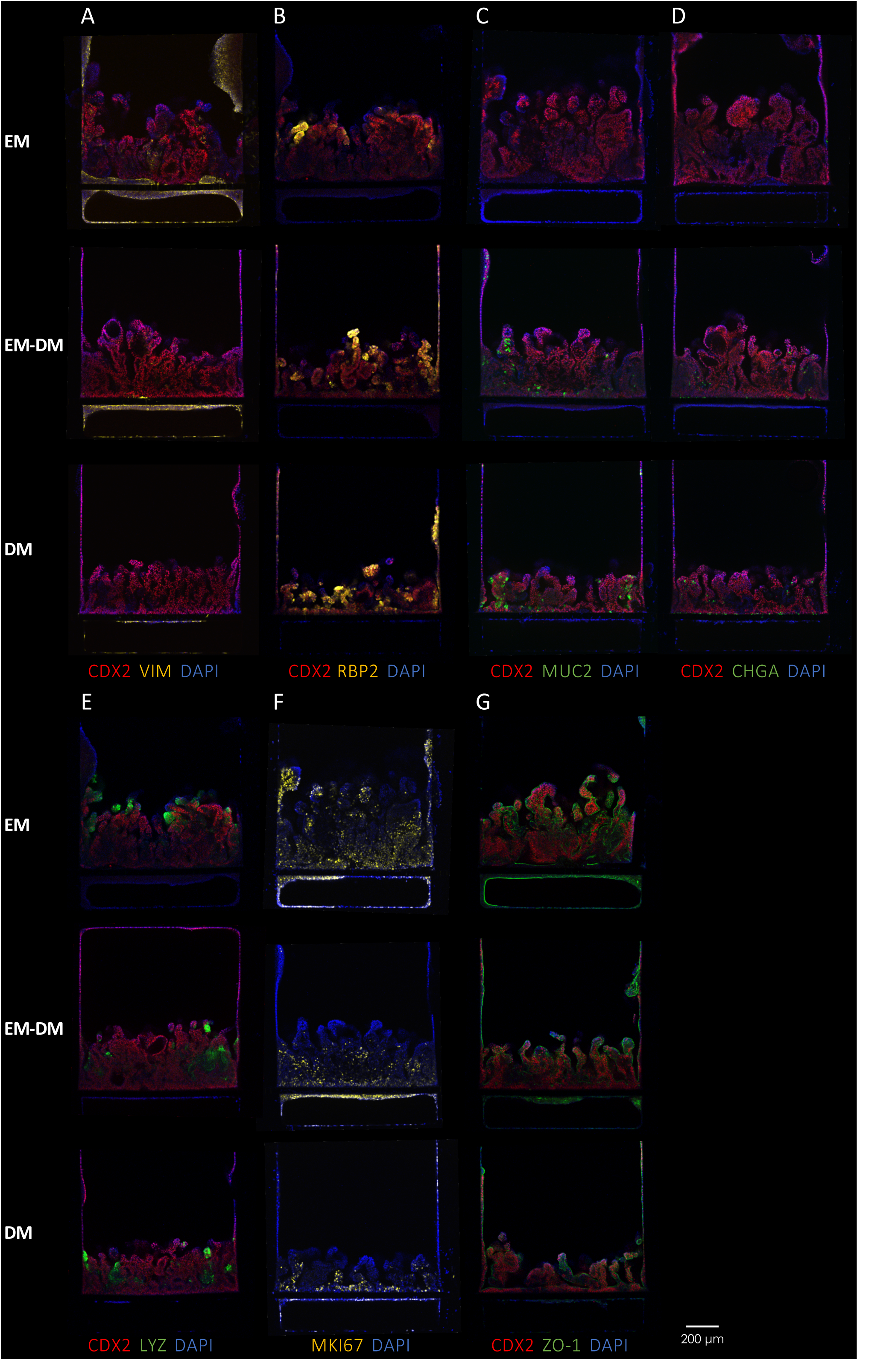
Diversification of epithelial cell-type composition upon exposure to growth factor gradients. Representative immunofluorescent confocal images of cross-sectional slices of the intestine-on-chip stained for markers characteristic of intestinal epithelial and mesenchymal cells: intestinal epithelial cells (CDX2), mesenchymal cells (VIM), enterocytes (RBP2), goblet cells (MUC2), enteroendocrine cells (CHGA), Paneth cells (LYZ), dividing cells (MKI67) and tight junctions (ZO-1).

The EM-DM and DM conditions efficiently induced the expression of markers associated with multiple mature intestinal epithelial subtypes such as enterocytes (RBP2-positive), goblet cells (MUC2-positive) and enteroendocrine cells (CHGA-positive), which were lowly expressed or absent in the EM condition (Figures 2B-D and S3D). Cells expressing lysozyme, a marker classically associated with Paneth cells, were detected in all conditions but were slightly enriched in the EM-DM and DM conditions (Figures 2E and S3D). Proliferative epithelial and mesenchymal cells (MKI67-positive) were decreased in the EM-DM condition and only minimally present in the DM condition when compared with the EM condition (Figure 2F). The localization of tight junction protein ZO-1 indicated polarization of the tissue with the apical side projected toward the top channel (Figure 2G). Thus, exposure to the EM-DM condition appears to preserve intestinal epithelial tissue morphology and proliferation while efficiently inducing mature epithelial subtypes and supporting the self-organization of mesenchymal cells in the bottom channel of the intestine-on-chip.

### Single-cell RNA-sequencing reveals the major intestinal epithelial subtypes, myofibroblasts and neurons

To obtain an untargeted high-resolution characterization of the cellular diversity in the hiPSC-derived intestine-on-chip in the EM, DM and EM-DM conditions, we performed scRNAseq on cells dissociated from the chips on day 6. In concordance with the microscopy and flow cytometry data, we identified two transcriptionally distinct cellular compartments: one corresponding to intestinal epithelial cells (*EPCAM-*, *CDX2-* and *CDH1*-positive) and an *EPCAM*-negative and *VIM*-positive compartment (Figures 3A, 3B and S4A). The latter included markers characteristic of both mesenchymal and neural cells, so we designated it the ‘mesenchymal/neural compartment’. In the epithelial compartment, we identified five main epithelial subtypes based on the concordance of differentially expressed genes with canonical markers^24–28^: transit-amplifying (TA)/stem cells (*SMOC2, LGR5, MKI67, TOP2A, PCNA* and *CENPF*), enterocytes (type 1: *FABP2, RBP2, APOA4, CYP3A5, ALDOB, ANPEP, SI* and *SLC2A2*; type 2: *SLC39A4, MT1E*, *MT1G, MT1H* and *MT2A*; progenitor: *FGB* and *FGG*), Paneth-like cells (*LYZ* and *PRSS2*), goblet cells (*MUC2, TFF3, ZG16* and *CLCA1*) and enteroendocrine cells (*CHGA, NEUROD1, MLN, GHRL* and *GCG*) (Figures 3A, 3B and S4B). Both type 1 and type 2 enterocytes expressed canonical enterocyte markers, but the type 2 enterocytes also expressed metallothioneins and metal transporters (e.g. zinc transporter *SLC39A4*), indicating a role in the metabolism of metals (Figures 3B and S4C). The Paneth-like cells expressed two Paneth-characteristic genes (*LYZ* and *PRSS2*) and showed enrichment for the antimicrobial humeral response pathway, but they did not express defensin genes and were therefore termed ‘Paneth-like cells’ (Figures 3B and S4C). Two epithelial clusters expressed both *EPCAM* and *VIM* and were therefore named ‘mesenchymal-like epithelial cell type 1 and 2’ (Figure 3B). A third epithelial cluster was only partly *VIM*-positive and was designated ‘mesenchymal-like epithelial precursor’ (Figures 3B and S4A). In particular, the mesenchymal-like epithelial cell type 2 cluster and, to a lesser extent, the precursor cluster shared many enriched pathways with the mesenchymal/neural compartment (Figure S4C). To simplify further analysis, we grouped enterocyte types 1 and 2 and enterocyte progenitors as ‘enterocytes’ and mesenchymal-like epithelial cell types 1 and 2 and precursor cells as ‘mesenchymal-like epithelial cells’.

**Figure 3.**
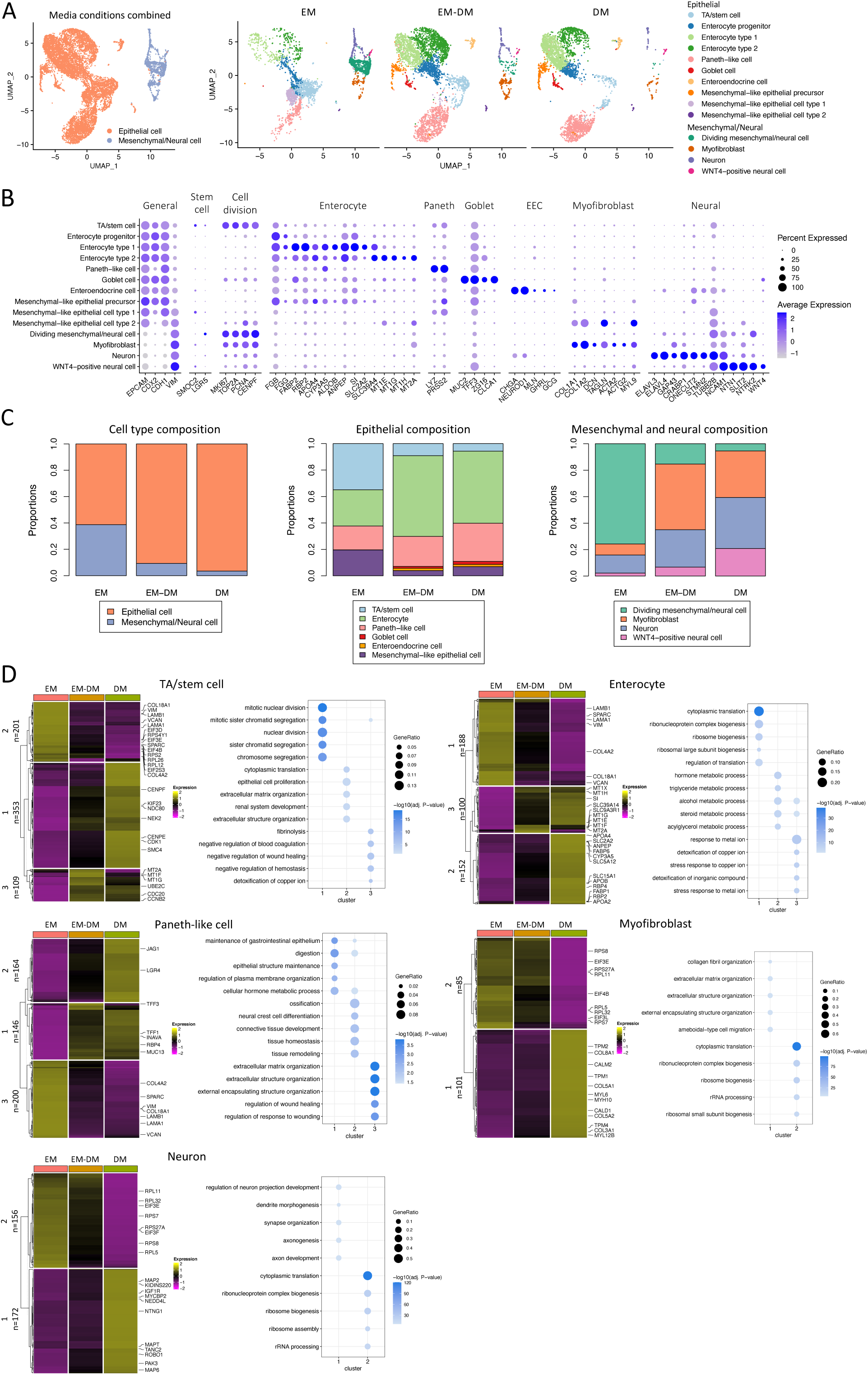
Characterization of cell-type composition and functionalities in different media conditions using single-cell RNA sequencing. (A) UMAP projection of the epithelial and mesenchymal/neural compartments (data of media conditions combined) and cell types. (B) Average expression of differentially expressed genes (DEGs) between cell types that correspond to canonical markers. The highest average log2 fold-change value for *LGR5* was 0.32 in dividing mesenchymal/neural cells (below the threshold of 0.5), but we show *LGR5* given its relevance for detecting intestinal stem cells. EEC = enteroendocrine cell. (C) Proportions of compartment and cell-type composition. (D) Heatmap of the average gene expression of the DEGs between media conditions, clustered in groups of genes with similar expression profiles. Clusters were used to perform pathway enrichment analysis using the Gene Ontology: biological process database. Relevant genes from the assigned pathways are highlighted in the heatmap. The number of genes assigned to each cluster (n) is indicated.

In the mesenchymal/neural compartment, we identified four clusters that expressed markers characteristic of the intestinal mesenchyme and enteric nervous system^26–28^: dividing mesenchymal/neural cells (*MKI67, TOP2A, PCNA* and *CENPF*), myofibroblasts (*COL1A1, COL1A2, DCN, TAGLN, ACTA2, ACTG2* and *MYL9*), neurons (*ELAVL3, ELAVL4, GAP43, CRABP1, ONECUT2, STMN2* and *TUBB2B*) and WNT4-positive neural cells (*NCAM1, NTN1, SLIT2, NTRK2* and *WNT4*) (Figures 3A, 3B and S4B). The myofibroblasts expressed fibroblast-characteristic genes related to collagen production, as well as smooth muscle-related actin genes and myosin gene *MYL9*, but lacked expression of the smooth-muscle-characteristic genes *DES* and *MYH11* (Figure 3B)^27^. Therefore, their expression profile corresponded most with myofibroblasts^23^. The WNT4-positive neural cells were enriched for genes related to axon guidance and expressed *WNT4*, which correlates with glial cell functions, but characteristic glial markers (*S100B*, *SOX10* and *PLP1*) were not expressed (Figure 3B)^26,27,29^. In concordance with the observation of a cluster expressing neuronal genes, we observed cells that morphologically resembled neurons in the bottom channel of the intestine-on-chip (Figure S5A). The emergence of neurons during the development of hiPSC-derived intestinal organoids was previously described to occur very rarely^15^. In our intestine-on-chip system, however, neurons developed consistently between biological replicates and in all media conditions (Figure S5B).

The ratio between the epithelial and mesenchymal/neural compartment and the subtype composition were reproducible between biological replicates, but varied greatly between the different media conditions (Figures 3C and S5B). Overall, the EM condition contained the largest proportion of mesenchymal/neural cells (38.7% (SD 10)) compared to the EM-DM (9.3% (SD 4.9)) and DM conditions (3.5% (SD 1.1)) (Figure 3C). In the EM condition, both the epithelial and mesenchymal/neural compartments contained a large proportion of dividing cells (epithelial: 42.5% (SD 4.4), mesenchymal/neural: 85.2% (0.7)) (Figures 3C and S5C). In the EM-DM and DM conditions, the frequency of dividing cells was reduced (epithelial: 16.3% (SD 5.8), 11.2% (SD 2); mesenchymal/neural: 59.6% (SD 7.9), 31.6% (SD 10.1), respectively), in line with the reduced tissue height and cell density in these conditions (Figures S1B, S1C and S5C). The reduction of dividing cells in the EM-DM and DM conditions corresponded predominantly with an increased frequency of enterocytes, goblet and enteroendocrine cells in the epithelial compartment and an increased frequency of myofibroblasts, neurons and WNT4-positive neural cells in the mesenchymal/neural compartment (Figure 3C, Table S1, S2).

These observations indicate that the intestine-on-chip exposed to the EM-DM condition captures and balances the dividing, progenitor and mature epithelial cell stages as well as dividing mesenchymal/neural cells, myofibroblast and neural populations.

### Cell types display different functionalities depending on media condition

To investigate the media-induced differences in epithelial, mesenchymal and neural subtypes, we performed differential gene expression and pathway enrichment analysis between the media conditions. We selected five subtypes to include in this analysis (TA/stem cells, enterocytes, Paneth-like cells, myofibroblasts and neurons) because they contained 30 or more cells in each media condition (Table S1). The TA/stem cells expressed cell-division-related genes regardless of the media condition, but processes related to different phases of the cell cycle were induced: genes involved in translation were upregulated in EM and genes involved in mitosis were upregulated in DM, which reflected the enrichment of TA/stem cells in the G2/M phase relative to the S phase in the EM-DM and DM conditions (Figures 3D and S5C). Enterocytes showed an upregulation of genes related to proliferation in the EM condition and to mature intestinal functions such as nutrient and drug metabolism and uptake (e.g. *SI, ANPEP, CYP3A5, SLC2A2, SLC5A12*) in the DM condition (Figure 3D)^25^. The enterocytes in the EM-DM condition displayed both proliferative and mature intestinal functions, reflecting the more balanced division between progenitor and mature stages (Figure 3D). The DM condition induced Paneth-like cells enriched for pathways involved in maintenance of the gastrointestinal epithelium and tissue homeostasis, including genes *INAVA, MUC13, RBP4, TFF1, TFF3* and Notch ligand *JAG1* and WNT receptor *LGR4* (Figure 3D). The myofibroblast and neuron populations in EM were enriched for translation and cell-division-related genes such as ribosomal proteins and translation initiation factors, while the DM condition induced genes related to extracellular matrix (ECM) organization (e.g. *COL3A1, COL5A1, COL5A2, COL8A1*) and muscle contraction (e.g. *MYL6, MYH10, MYL12B, CALD1, CALM2* and *TPM1/2/4*) in myofibroblasts and genes related to axon, dendrite and synapse development and organization in neurons (Figure 3D). Overall, the EM condition induced proliferation-related genes and pathways in the epithelial, mesenchymal and neural subtypes in the intestine-on-chip, whereas the DM condition induced mature functionalities. Cell types exposed to the EM-DM condition displayed characteristics of both, demonstrating their intermediate or more heterogeneous state relative to the EM and DM conditions.

### The intestine-on-chip shows resemblance to the human small intestine

To investigate whether the hiPSC-derived intestine-on-chip is a representative model of the human small intestine, we compared our dataset to scRNAseq data of human intestinal tissue. Using the Human Fetal Endoderm Atlas^30^, we confirmed the overall intestinal phenotype of the hiPSC-derived intestine-on-chip relative to other organs derived from the endoderm germlayer (Figure S6A). Using the reference dataset from the Gut Cell Atlas^27,31^, we predicted the age, intestinal region and cell-type identity of the cells in the intestine-on-chip (Figure 4A-C). The majority of cells in the intestine-on-chip system were predicted to resemble an early developmental fetal stage of the human intestine corresponding to ‘first trimester’ (Figure 4A). Only the most mature enterocytes and Paneth-like cells were projected as ‘second trimester’ and ‘adult’, respectively. The intestinal region was projected to be ‘small intestine’ for the majority of cells in the EM-DM and DM conditions, while half of the cells were projected as ‘large intestine’ in the EM condition (Figure 4B).

**Figure 4.**
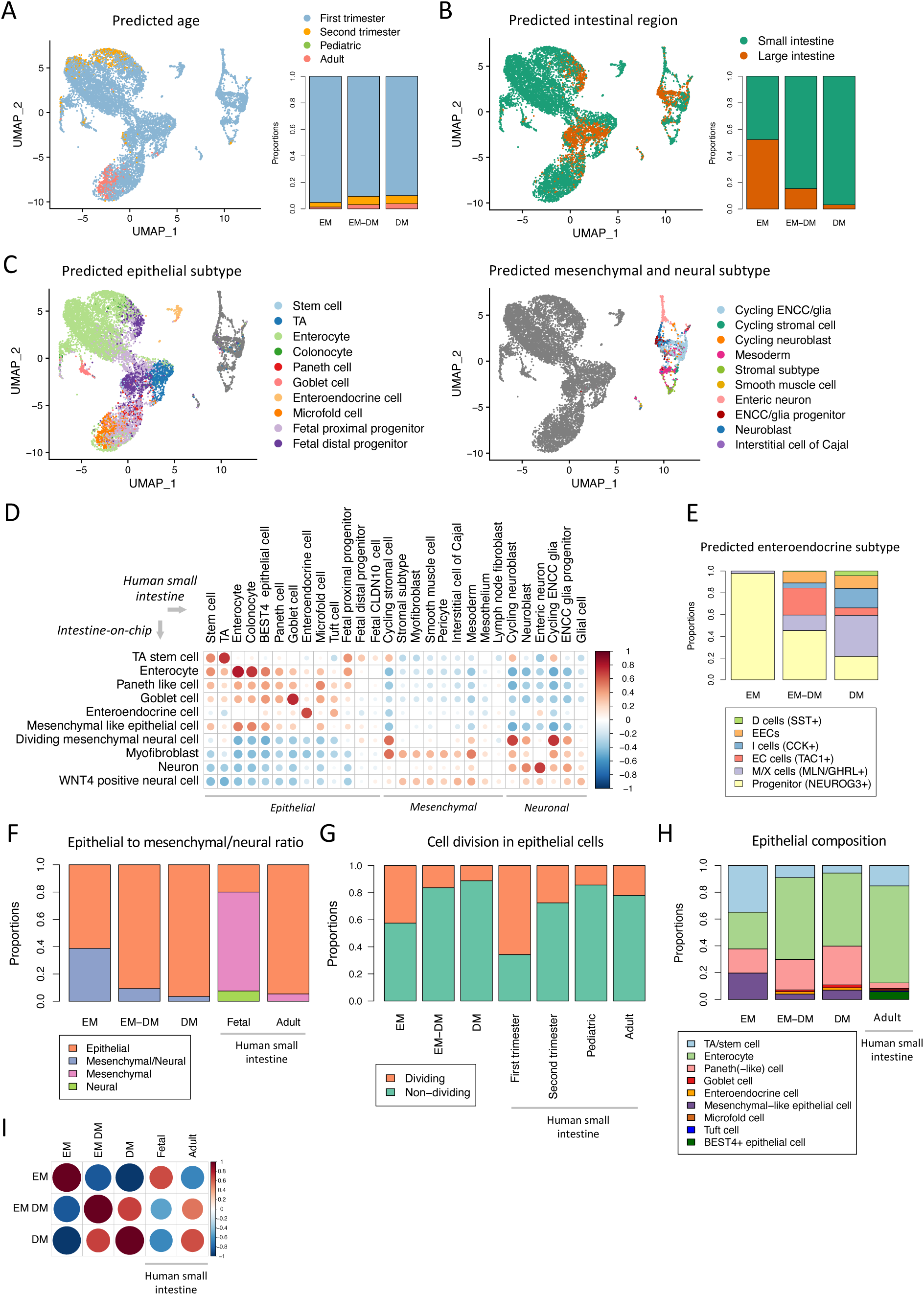
Comparison of the intestine-on-chip to the human intestine. UMAP projection and proportions of predicted age group (A), intestinal region (B) and cell types (C) based on human intestinal data of the Gut Cell Atlas (including the age groups first trimester, second trimester, pediatric and adult; the regions small intestine, large intestine, appendix, rectum and mesenteric lymph node; and the compartments epithelial, mesenchymal, neural, endothelial and immune cells). Predicted cell types with >10 cells are shown. (D) Correlation analysis between cell types in the intestine-on-chip and the epithelial, mesenchymal and neural subtypes in the human small intestine from all age groups (Gut Cell Atlas). Correlation scores were generated based on the average expression of differentially expressed genes (DEGs) between the cell types in the intestine-on-chip. Color intensity and dot size correspond to Pearson correlation scores and non-significant correlations (p-value > 0.01) were left blank. (E) Proportion of predicted types of enteroendocrine cells in the intestine-on-chip based on human intestinal data of the Gut Cell Atlas as described for (C). (F) Proportion of epithelial, mesenchymal and neural cells in the intestine-on-chip and the human fetal and adult small intestine (Gut Cell Atlas). (G) Proportion of dividing and non-dividing epithelial cells in the intestine-on-chip and the human fetal, pediatric and adult small intestine (Gut Cell Atlas). (H) Proportion of epithelial subtypes in the intestine-on-chip and the human adult small intestine (Gut Cell Atlas). (I) Correlation analysis between media conditions in the intestine-on-chip and epithelial, mesenchymal and neural cells in the human fetal and adult small intestine (Gut Cell Atlas). Correlation scores were generated based on the average expression of DEGs between the media conditions in the intestine-on-chip. Color intensity and dot size correspond to Pearson correlation scores and non-significant correlations (p-value > 0.01) were left blank.

Our cell-type cluster assignments were validated by projecting the cells of the intestine-on-chip onto human intestinal cell clusters, with a few notable differences. The Paneth-like cell population was partly predicted as Paneth cells, but also as microfold and fetal proximal progenitor cells, the dividing mesenchymal/neural cells were projected as neural cells (predominantly ‘cycling enteric neural crest cells/glia’) rather than mesenchymal cells, and the WNT4-positive neural cells were mainly projected as ‘enteric neural crest cell/glia progenitor’ (Figures 4C and S6B). These results were verified by correlation analysis, which demonstrated a high correlation between the cell types from the intestine-on-chip and the respective cell types in the human small intestine, except for the Paneth-like cells, which did not strongly correlate to a specific epithelial subtype (Figure 4D). Additionally, the validation of cell-type identities using reference data provided deeper insight into the predicted subtypes of enteroendocrine cells in our intestine-on-chip system. In the EM-DM and DM conditions, cells resembling M/X, I, enterochromaffin and D cells were identified (Figure 4E). The expression of hormones corresponding to these and several other subtypes (L, N and S cells) was confirmed in the scRNAseq data of the enteroendocrine population in the intestine-on-chip (Figure S6C)^32^.

Finally, we compared the cell-type ratios of the intestine-on-chip to the ratios as described for the human small intestine by the Gut Cell Atlas dataset. In the EM condition, the epithelial to mesenchymal/neural ratio and the proportion of dividing epithelial cells most resembles the composition of the fetal human small intestine, while in the EM-DM and DM conditions the composition shifts toward that found in the human adult small intestine (Figures 4F and 4G). In addition, the epithelial subtype composition shifts toward one that resembles the human adult small intestine, with the EM-DM condition showing the closest approximation of the three media conditions (Figure 4H, Table S2). The DM-induced maturation of the cells in the intestine-on-chip toward a later developmental state was underlined by correlation analysis using the differentially expressed genes between media conditions (Figure 4I).

Overall, the intestine-on-chip was predicted to have a small intestinal and fetal phenotype in the EM, EM-DM and DM conditions, when compared to the human intestine. However, we can induce maturation and steer the system toward a cell-type composition that resembles the adult human intestine by exposure to the EM-DM condition.

### Identification of differentiation trajectories and processes driving lineage induction

Intestinal epithelial differentiation progresses along the crypt–villus axis from stem cells and TA cells toward mature absorptive and secretory cell types. The lineage specification is regulated by timely induction or repression of several transcription factors^1^. We performed trajectory and pseudotime analysis to investigate whether the differentiation patterns of intestinal epithelial lineages in the intestine-on-chip follow patterns similar to those described for the human intestine. The datasets of all three media conditions were combined to infer overall directionality in the intestine-on-chip data using RNA velocity. For both the epithelial and mesenchymal/neural populations, differentiation trajectories and pseudotime analysis demonstrated directionality from dividing cells (TA/stem cells and dividing mesenchymal/neural cells, respectively) to mature cell types (Figures 5A, 5B and S7A). Interestingly, the neuronal and myofibroblast populations were predicted to emerge from the same population of dividing mesenchymal/neural cells (Figure 5A). To further investigate the mechanism underlying lineage induction, we identified putative driver genes involved in fate specification of enterocytes (type 1 and type 2), Paneth-like cells, enteroendocrine cells, myofibroblasts and neurons (Figures 5C, S7B). Previously described driver genes for the epithelial lineages were highlighted and were especially prevalent in the enteroendocrine lineage (i.e. *NEUROG3, PAX4, NKX2.2, FEV*) (marked by arrows in Figure 5C)^1,25,33–39^. Together, this data provides insight into the differentiation trajectories of hiPSC-derived intestinal lineages in the intestine-on-chip and indicates similarity to those described for human or murine adult stem cell-derived intestinal tissue.

**Figure 5.**
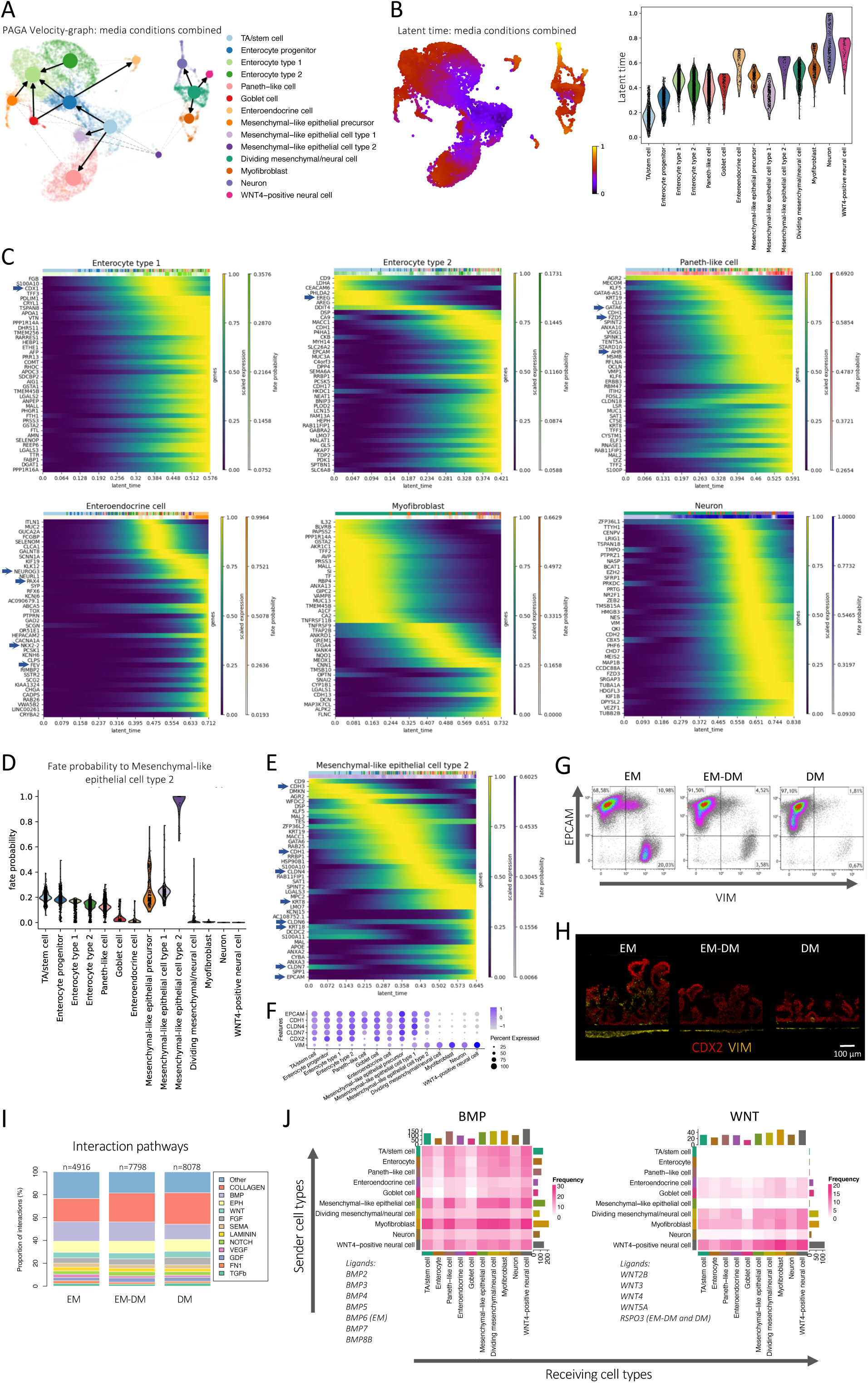
Characterization of differentiation trajectories and cell–cell communication. (A) Differentiation trajectories of cell types in the intestine-on-chip projected by Partition-based Graph Abstraction (PAGA) analysis extended by RNA velocity-inferred directionality (scVelo). (B) Latent time projections derived from RNA velocity analysis (scVelo). (C) Temporal activation of the top putative driver genes along specific trajectories determined by fate probability analysis (scVelo and CellRank). Putative driver genes that correspond to previously reported driver genes in adult stem cell-derived intestinal tissue are marked by arrows. (D) Fate probability to contribute to the differentiation lineage toward the mesenchymal-like epithelial cell type 2 cluster (scVelo and CellRank). (E) Temporal activation of the top putative driver genes along the trajectory of mesenchymal-like epithelial cell type 2 determined by fate probability analysis (scVelo and CellRank). Putative driver genes that are associated with epithelial-to-mesenchymal transition are marked by arrows. (F) Gene expression of characteristic intestinal epithelial markers (*EPCAM*, *CDX2* and junction markers: *CDH1, CLDN4* and *CLDN7*) and mesenchymal-marker *VIM*. (G) Protein expression of EPCAM and VIM determined by flow cytometry. (H) Protein expression of CDX2 and VIM determined by microscopy. Representative immunofluorescent confocal images of cross-sectional slices of the intestine-on-chip are shown. (I) Proportions of potential cell–cell interactions between cell types, grouped by signaling pathways, predicted by ligand–receptor expression analysis (Liana, CellPhoneDB). The pathways with a contribution lower than 2% in all media conditions were grouped as ‘other’. The number of interactions predicted in each media condition (n) is indicated. (J) Frequency of potential communication events between cell types for the BMP and WNT (canonical and non-canonical) pathways determined by ligand–receptor expression analysis (Liana, CellPhoneDB). The analysis was performed using the combined dataset of the media conditions. The detected ligands for each pathway are shown. BMP6 was only detected as ligand in the EM condition and R-spondin 3 was only detected in the EM-DM and DM conditions.

### Epithelial-to-mesenchymal transition in the EM condition

In the epithelial compartment of the intestine-on-chip, we identified several cell clusters (mesenchymal-like epithelial precursor and cell types 1 and 2) that displayed characteristics of both epithelial and mesenchymal cells as they expressed *EPCAM* and *VIM* and/or were enriched for pathways shared with the mesenchymal/neural compartment (Figures 3B and S4C)^40^. We investigated whether these cell clusters were indicative of epithelial-to-mesenchymal transition (EMT) by predicting which cells in the intestine-on-chip were likely to contribute to a certain differentiation lineage and hereby infer cell fate. These ‘fate probabilities’, together with trajectory analysis, predicted that some epithelial subtypes, but especially the mesenchymal-like epithelial precursors and type 1 cells had an increased probability to contribute to the mesenchymal-like epithelial cell type 2 cluster (Figure 5D) and that differentiation to this cluster included many putative driver genes related to EMT, including claudins, cadherins and cytokeratins (marked by arrows in Figure 5E)^41^. The expression of several of these epithelial-characteristic junction genes was reduced in the type 2 cluster relative to other epithelial clusters, indicating the EMT-associated loss of polarity (Figure 5F)^40,41^. Additionally, the type 2 cluster expressed genes that were involved in the same pathways that were enriched in the myofibroblast cluster, including many pathways related to ECM organization (Figure S4C). The presence of an EPCAM and VIM double-positive population was verified by flow cytometry data, indicating a high abundance of double-positive cells in the EM condition compared to the EM-DM and DM conditions (Figures 5G). In concordance, CDX2 and VIM double-positive cells were observed in the microscopy data of the EM condition (Figure 5H). These observations are in line with the scRNAseq data, which demonstrated a high proportion of mesenchymal-like epithelial cells in the EM condition (Figure 3C). Not only the mesenchymal-like epithelial clusters displayed EMT signatures: the same set of EMT-related genes (*VIM, COL4A2, COL18A1, SPARC, LAMA1, LAMB1* and *VCAN*) was differentially expressed in TA/stem cells, enterocytes and Paneth-like cells in the EM condition compared to the EM-DM and DM conditions (Figure 3D)^42^. Accordingly, pathways related to ECM organization were enriched in the TA/stem cell and Paneth-like cell clusters in the EM condition (Figure 3D). Together, this data indicates a transition from epithelial subtypes to an EPCAM- and VIM-positive state and then to a fibroblast-like phenotype, predominantly in the EM condition. EMT is an essential and abundant process during embryonic development^41^, which suggests that the EM condition may resemble a more embryonic growth environment that emphasizes the fetal character of hiPSC-derived tissues.

### Potential communication between intestinal epithelial, mesenchymal and neural cells

Intestinal epithelial fate specification and differentiation requires local activation and inhibition of the WNT and BMP pathways, which is facilitated by signals from the microenvironment, specifically the underlying mesenchymal compartment^1^. We investigated cell–cell communication between epithelial, mesenchymal and neural subtypes in the intestine-on-chip by studying potential interactions through the expression of ligand– receptor pairs. The BMP and WNT pathway were among the pathways that showed the most potential interactions in the intestine-on-chip (Figure 5I). The myofibroblast and WNT4- positive neural cell populations were predicted to be the most potent senders of BMP and WNT ligands, including BMP2 and BMP4 (the main BMPs in the human intestine^1^) and WNT2B, WNT3, WNT5A and R-spondin 3, a potentiator of WNT signaling that was exclusively detected as a ligand in the EM-DM and DM conditions (Figure 5J). In the intestine-on-chip, the BMP pathway was inhibited by supplementation of Noggin in EM, but not in DM, and no external BMP ligands were included in either medium. This allows for potential effects from intrinsically secreted BMP ligands in the EM-DM and DM conditions. The WNT pathway is activated in EM by the inclusion of CHIR99021, which might make cell-derived WNT ligands redundant, but DM does not include external WNT ligands and might benefit intrinsically induced WNT signaling. Lastly, many ligand–receptor interactions were predicted through ECM components (e.g. collagen and laminin) in the intestine-on-chip (Figures 5I and S7C). ECM–cell interactions are essential in tissue remodeling and organization and cell differentiation processes in the human intestine^43^ and might contribute to the self-organization of the cells in the intestine-on-chip. Similarly, the ephrin and semaphorin pathways, which were identified to contain many interactions within the intestine-on-chip, are described to be involved in tissue organization and cell guidance (Figures 5I and S7C)^44,45^. Overall, mesenchymal and neural subtypes in the intestine-on-chip express ligands that could support epithelial cell fate to mature subtypes, the maintenance of stem cell function and tissue organization. However, the relevance of these potential interactions in the intestine-on-chip still needs to be addressed, as several of these pathways are also activated or inhibited via factors in the medium.

### Selective sensitivity of cell types to type I or type II interferon stimulation

To test the potential of the intestine-on-chip system for studying inflammation in the context of intestinal diseases, we exposed the system to type I and type II interferons (IFNs), pro-inflammatory cytokines involved in multiple intestinal diseases^46,47^. IFN-β (500 U/ml) or IFN-γ (100 ng/ml) was added to the bottom channel of the intestine-on-chip in the EM-DM condition for 6 hours to simulate an inflammatory state in the lamina propria of the human intestine, and scRNAseq was performed. The responsiveness of the cells in the intestine-on-chip to both types of IFNs was verified by differential gene expression and pathway enrichment analysis (Figures 6A and S8A). To investigate differences in the sensitivity of the cells in the intestine-on-chip to IFN stimulation, IFN-β-responding cells (*IRF1-, APOL6- and TAP1*-positive) and IFN-γ-responding cells (*IFI6-, IFIT1- and MX1-*positive) were annotated (Figures 6B and S8B). The genes used for annotation are known type I and type II IFN response genes^48^ that were differentially expressed in both the epithelial and mesenchymal/neural compartments of the intestine-on-chip upon IFN stimulation (Figure S8A). Interestingly, some but not all cells in the intestine-on-chip responded to these cytokines, even though the IFN-β and IFN-γ receptors were homogenously expressed in the epithelial, mesenchymal and neural subtypes (Figures 6B-D). Strikingly, only 10.3% (SD 0.02) of the epithelial population responded to IFN-β and 30.6% (SD 0.05) to IFN-γ, while 68.2% (SD 0.09) of the mesenchymal/neural population responded to IFN-β and 66.8% (SD 0.03) to IFN-γ (Figure 6D). The difference in IFN sensitivity between epithelial cells and mesenchymal/neural cells might be explained by the fact that the mesenchymal and neural cells were in direct contact with the cytokines in the bottom channel of the intestine-on-chip (Figure 2A).

**Figure 6.**
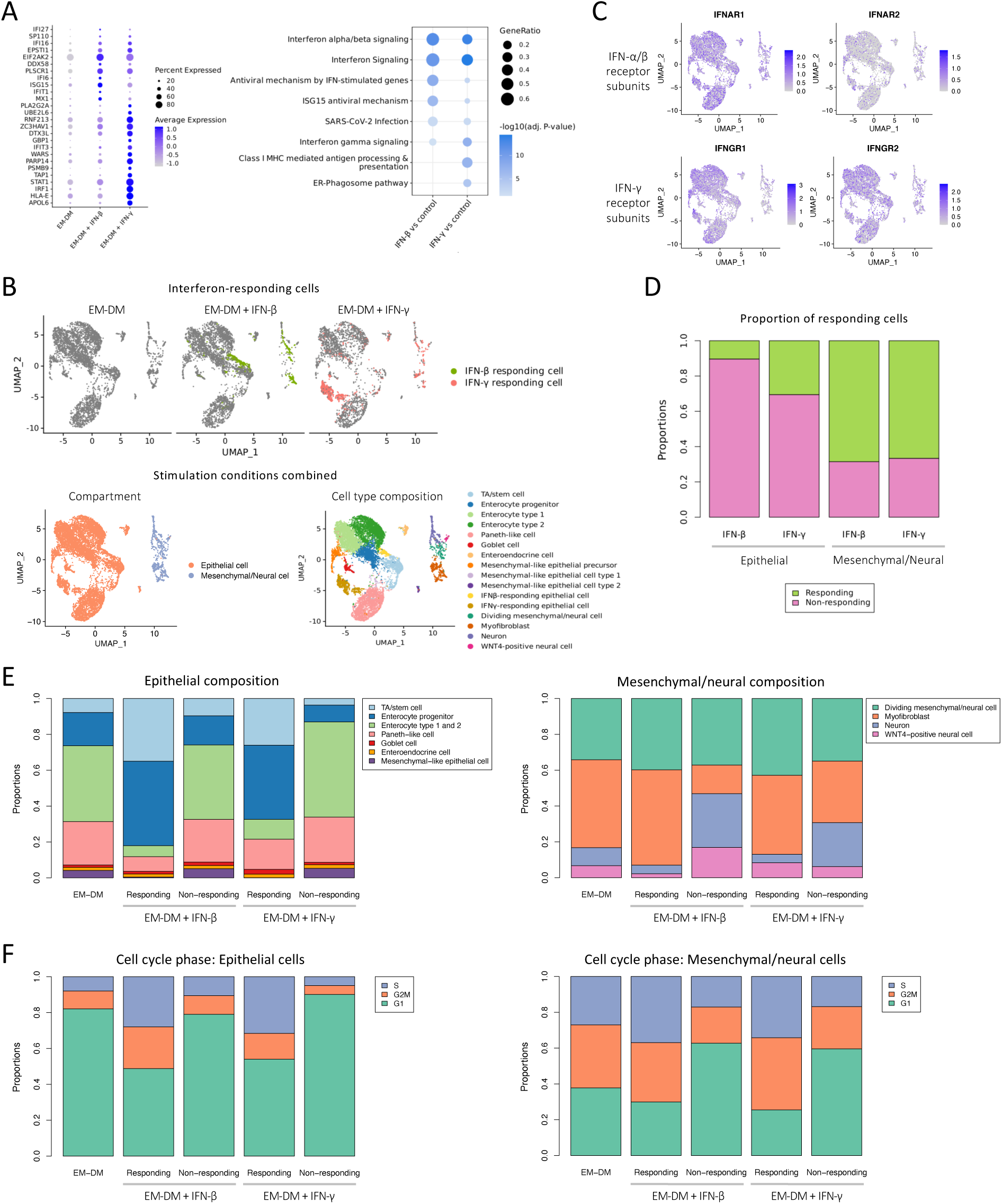
Selective cellular response of epithelial, mesenchymal and neural cells to IFN-β and IFN-γ. (A) Average expression of the top differentially expressed genes (DEGs) between the EM-DM and interferon (IFN)-β-stimulated or IFN-γ-stimulated condition. All DEGs were used for pathway enrichment analysis using the Reactome database. (B) UMAP projection of IFN-β and IFN-γ responding cells in the individual datasets and the cellular compartment and cell-type annotation in the combined dataset of the EM-DM, IFN-β-stimulated and IFN-γ-stimulated conditions. (C) UMAP projection of expression of type I IFN (*IFNAR1* and *IFNAR2*) and type II IFN (*IFNGR1* and *IFNGR2*) receptor subunits using the combined dataset of the EM-DM, IFN-β-stimulated and IFN-γ-stimulated conditions. (D) Proportion of IFN-responding epithelial and mesenchymal/neural cells after IFN-β or IFN-γ stimulation. (E) Proportion of epithelial, mesenchymal and neural subtypes in IFN-responding and non-responding cells. (F) Proportion of cells in a specific cell-cycle phase in the IFN-responding and non-responding epithelial, mesenchymal and neural cells.

To identify whether specific epithelial, mesenchymal and neural subtypes responded to IFN stimulation, we integrated the datasets of the unstimulated EM-DM, IFN-β-stimulated and IFN-γ-stimulated conditions while correcting for cytokine stimulation and investigated which cell types most resembled the IFN-responding cells (Figure S8C). We identified an enrichment of TA/stem cells and enterocyte progenitors in the IFN-responding epithelial cells and an enrichment of dividing mesenchymal/neural cells and myofibroblasts in the IFN- responding mesenchymal/neural cells (Figure 6E). This aligns with the observation that a high percentage of the IFN-β- and IFN-γ-responding cells were dividing cells (e.g. cells in the S and G2/M phase of the cell cycle) when compared to non-responding cells (Figure 6F). Indeed, the type I IFN response has been shown to be dependent on cell-cycle state at single-cell level^49^, so cell-cycle state may explain the selective response of cell types to IFNs we observe as we only stimulated the intestine-on-chip for 6 hours.

The hiPSC-derived intestine-on-chip can be used to model selective cellular responses to external factors, such as cytokines. Dividing epithelial, mesenchymal and neural subtypes were shown to be more responsive to IFN-β and IFN-γ stimulation than non-dividing cells indicating differences in IFN sensitivities between cells in the intestine-on-chip.

## Discussion

### Main outcome

We developed a hiPSC-derived intestine-on-chip, which includes a diverse epithelial composition and myofibroblast and neural subtypes, as a model for the human small intestine. Exposure to EM in the bottom channel and DM in the top channel of the system, replicating the growth factor gradients in the human intestine, induced an intestinal epithelial subtype composition that balances proliferating and mature secretory and absorptive subtypes and reflects the human adult small intestine. The epithelial cells, which self-organized into villus-like folds in the top channel, and the myofibroblast and neural cell types, which emerged in the bottom channel, could be sustained in a spatially confined manner for (at least) 6 days. Conversely, exposure to EM in both compartments induced an over-representation of mesenchymal and neural cells and dividing epithelial cells with hallmarks of EMT, while exposure to DM in both compartments induced an enrichment for mature epithelial subtypes and preserved very few mesenchymal, neural cells and dividing epithelial cells resulting in greatly shortened villus-like folds. In summary, combined exposure to EM and DM in a hiPSC-derived intestine-on-chip system provides a sustainable and reproducible method to model the human small intestinal epithelial barrier, with physiologically relevant cellular diversity, including myofibroblast and neural cell types.

### Media composition

The composition of EM and DM included compounds that activate or inhibit well-established pathways in intestinal development and were previously demonstrated to be effective in controlling epithelial lineage induction in adult stem cell-derived intestinal epithelial organoids^50,51^. Overall, we have demonstrated that the same differentiation factors used in adult stem cell-derived intestinal organoids can efficiently induce mature lineages in hiPSC-derived intestinal epithelial cells, and even in mesenchymal and neural cells. Several putative driver genes identified for the epithelial lineages in this study were shown to correspond to established driver genes known from adult stem cell-derived intestinal tissue, suggesting that similar pathways are involved in lineage differentiation regardless of the different origins of the intestinal tissues.

### Cell-type composition

The gene expression profiles of all the epithelial subtypes observed in the EM-DM exposed intestine-on-chip, except for the Paneth-like cells, correlated highly and specifically to the profiles of their human *in vivo* counterparts. We identified two types of mature enterocytes in our system, designated type 1 and type 2 enterocytes. Type 1 enterocytes expressed genes representing enterocyte-characteristic functions in metabolism and uptake of lipids, peptides, carbohydrates, vitamins and drugs. Type 2 enterocytes were characterized by a strong upregulation of metallothioneins, which play a role in the absorbance of minerals and prevention of metal toxicity^52^. The enteroendocrine population expressed various hormones that represent different lineages (mainly M, X, S, N and L cells) characteristic of the intestinal crypt and villus region^53^. The Paneth-like cells in our system exhibited high expression of the Paneth-cell-characteristic genes *LYZ* and *PRSS2*^24,26^, but lacked expression of defensin genes, which might indicate limited maturity of these cells. Gene expression profiles in Paneth-like cells were not correlated to a specific human intestinal epithelial subtype, but indicated a resemblance to both microfold and Paneth cells, which could (partly) be explained by the shared expression of lysozyme in these two cell types^24^. However, no other microfold-cell-characteristic genes were identified in our data. The Paneth-like cell cluster had a high abundance in all media conditions, even though Paneth cells in the human intestine require active WNT signaling^1^, which was not induced in the DM condition. Moreover, the expression of lysozyme has been observed in duodenal organoids derived from fetal tissue taken before the onset of Paneth cells in the human intestine, which might underline the need for further maturation of Paneth-like cells in the intestine-on-chip^28^. Further Paneth cell induction and maturation might require exposure to additional growth factors, such as IL-22^54^. In addition to the absence of microfold cells, no Tuft cells were identified in the intestine-on-chip. Efficient induction of microfold cells and Tuft cells could potentially be achieved by the addition of RANKL and a combination of IL-4 and IL-13 to DM, respectively^55,56^.

hiPSC-derived mesenchymal and neural subtypes co-differentiated with epithelial cells when exposed to EM-DM or DM and included populations resembling myofibroblasts and neurons. The gene expression profile of the neurons in the intestine-on-chip strongly correlated with neurons of the human enteric nervous system and included canonical marker genes (e.g. *ELAVL3, ELAVL4* and *TUBB2B*)^26,27^. Previous efforts to include neuronal subtypes in hiPSC-derived intestinal organoid cultures required a separate *in vitro* differentiation trajectory to generate enteric nervous system precursors, which were later combined with intestinal organoids^15^, or the substitution of EGF with EREG, which was not included here^57^. The co-development of neuronal cells in the intestine-on-chip, which was consistent between biological replicates and was observed in all media conditions, might thus be the result of the dynamic forces the cells are exposed to in this system when compared to conventional static cultures. The myofibroblast cells displayed a high expression of ECM proteins, such as collagen, and several genes related to muscle contraction. The co-development of (myo)fibroblast and smooth muscle cells in hiPSC-derived intestinal organoids *in vitro* was described earlier, however desmin-positive smooth muscle cells were not observed in our system^21^. The WNT4-positive neural cell population found in the intestine-on-chip, was difficult to correlate to a specific mesenchymal or neural human subtype, but it seemed specifically functional in axon guidance and support of the epithelial layer and might be glial progenitor cells^29^.

The co-development of mesenchymal cells during the *in vitro* differentiation of hiPSCs toward intestinal epithelial cells was previously attributed to the development of mesoderm precursors during endoderm induction in the initial stages of differentiation^21^. Our data suggest that mesenchymal cells might also develop in later or final stages of this differentiation due to induction of EMT, particularly in the EM condition. The reduced abundance of mesenchymal-like epithelial cells in the EM-DM and DM conditions might be related to the inhibition of the MAPK pathway, which was postulated as a mediator of EMT^58^. The myofibroblast and neural populations in the intestine-on-chip seemed to emerge from the same population of cycling cells with characteristics of mesenchymal and neural stem cells. The origin of the enteric nervous system in the human intestine has been debated in recent years, and it has been speculated that some neurons in the human intestine are not derived from neural crest cells but are rather of mesodermal and/or endodermal origin^59,60^. Nevertheless, the underlying mechanism of the development of myofibroblasts and neurons in the intestine-on-chip requires further investigation.

### Spatial organization

The spatial organization of intestinal epithelial, mesenchymal and neural cells in the intestine-on-chip partly captures the architecture of the human intestinal barrier. The epithelial cells form villus-like folds that are polarized, with the apical side, indicated by local zonulin-1 expression, facing the top channel. The mesenchymal and neural cells self-organize to form a layer underneath the epithelial compartment in the bottom channel, where they selectively and consistently emerge during the first 7 days after seeding in the top channel. These cells may simply migrate through the pores in the membrane between the two channels in search of space to populate, or they may be attracted by basolateral expression of chemoattractant molecules by the epithelial layer. The epithelial cells self-organize into villus-like folds exclusively and specifically on the membrane area directly above the bottom channel, in all media conditions. This suggests that the formation of villus-like folds depends on the continuous flow of medium on the basolateral side of the tissue, for example for the local removal of WNT antagonists (as has been shown earlier^61,62^), or the exposure to shear stress basolaterally. A current limitation of the intestine-on-chip is lack of physiological organization of the intestinal epithelial subtypes along these villus-like folds. This could be improved by providing a scaffold with crypt–villus structure. Such a scaffold was demonstrated to induce local stem and Paneth cell formation in the crypt region^63,64^. However, this requires a more complex system that might limit direct access to the basolateral side of the epithelial barrier.

### Resemblance to the human intestine

Comparison of the transcriptional profile of the intestine-on-chip with the human intestine in different developmental stages indicated that the intestine-on-chip resembles the fetal intestine more than the adult human intestine, in line with literature indicating the overall fetal phenotype of hiPSC-derived intestinal tissues^16^. Additionally, the high plasticity of the cells in the intestine-on-chip, which was demonstrated by the emergence of myofibroblasts and neurons seemingly from the same cycling cell population and the hallmarks of EMT, might indicate the relative fetal state of these hiPSC-derived tissues^41^. Maturation of hiPSC-derived intestinal organoids has previously been achieved by transplantation *in vivo* (in mice)^11,14–16^; however, this limits downstream applications. We demonstrate that exposure to a more physiologically relevant microenvironment can facilitate *in vitro* maturation of hiPSC-derived intestinal tissue. In particular, while the cell-type composition in the EM condition resembled the fetal small intestine, exposure to the EM-DM condition yielded compositions that resembled the adult human small intestine and induced gene expression profiles associated with mature functionalities in the different cell types (e.g. nutrient and drug metabolizing enzymes and transporters in enterocytes). Moreover, EMT-associated cell-type clusters and expression profiles were greatly reduced in the epithelial cells exposed to the EM-DM condition relative to the EM condition.

### Response to interferons

IFNs are implicated in multiple intestinal diseases, e.g. inflammatory bowel disease and celiac disease^46,47^, so the responsiveness of the cells in the intestine-on-chip to external factors such as IFNs is important for its applicability in disease modeling. Epithelial, mesenchymal and neural cells were shown to respond to IFN stimulation and expressed established type I or II IFN-inducible genes depending on the exposure to IFN-β or IFN-γ. However, the dividing epithelial, mesenchymal and neural cells in the intestine-on-chip showed an increased sensitivity to IFN-β and IFN-γ compared to non-dividing cells, despite a uniform expression of

IFN receptor genes. This observation is in line with a study that demonstrated cell-cycle-dependent responses to type I IFN at single-cell level^49^ and might relate to a study indicating that IFN-γ specifically targets intestinal stem cells in both human and mouse intestinal organoids^65^. As the IFN stimulation in the intestine-on-chip was only 6 hours, it is likely that only the cells in S and G2/M phase responded. The cell cycle is known to greatly influence gene expression profiles^66^, and certain activators or suppressors of IFN signaling may be present only in the S and/or G2/M phase, driving cell-cycle-dependent responses.

### Conclusion and future outlook

Dual exposure to expansion and differentiation media in a hiPSC-derived intestine-on-chip induces an epithelial cell-type composition reflective of the human adult small intestine and the presence of myofibroblasts and neurons. These human intestinal model systems with a more physiological microenvironment and increased complexity are valuable for disease modeling and studying the crosstalk between different subtypes. The iPSC basis of the model allows for personalization and establishment of reproducible disease models of, for example, inflammatory bowel disease or celiac disease patients, allowing for investigation of barrier function in the context of disease. Moreover, we envision that this system, which harbors different compartments, could support integration of additional cell lineages, e.g. mucosal immune cells, endothelial cells and microbial cells, or external factors such as complex mixtures of (drug) metabolites and nutritional compounds.

## Materials and methods

### Cell lines and culturing conditions

Three human induced pluripotent stem cell (hiPSC) lines were generated from renal epithelial cells derived from urine samples of three healthy donors (two male, one female) by the iPSC/CRISPR facility (ERIBA, UMCG, Groningen) using a lentiviral vector described earlier^67^. The expression of pluripotency markers and lack of differentiation markers in the resulting hiPSC lines was verified on protein and RNA level. The hiPSC lines were maintained in mTeSR Plus (Stemcell Technologies, #05825) on plates coated with hESC-qualified Matrigel (Corning #354277). Passaging was performed every 3-4 days using ReLeSR (Stemcell Technologies, #05872) according to manufacturer’s instructions. Cells were cultured in a humidified environment, at 37 °C in 5% CO2. hiPSC lines were cryopreserved as fragments in CryoStor CS10 (Stemcell Technologies #7930). All experiments with hiPSC lines were approved by the ethics committee of the University Medical Centre Groningen, document no. METC 2013/440 and written consent was obtained from all donors.

### Differentiation of hiPSCs to intestinal organoids

The differentiation procedure to generate intestinal organoids from hiPSCs was adapted from previously described protocols^17,18^. Briefly, hiPSCs were grown to ∼75-80% confluence in a 24-well plate. Definitive endoderm was induced by 3-day exposure to RPMI 1640 (ThermoFisher #11875093) supplemented with L-glutamine (2 mM, ThermoFisher #25030081), Penicillin-Streptomycin (100 Units/ml; 100 μg/ml respectively, ThermoFisher # 15140122), Activin A (100 ng/ml, R&D Systems, #338-AC) and an increasing concentration of defined FBS (day 1: 0%, day 2: 0.2%, day 3: 2% vol/vol, GElifesciences, Hyclone #SH30070.02). Additionally, Wnt3a was supplemented only on day 1 (25 ng/ml, R&D Systems, #5036-WN/CF). Thereafter, mid/hindgut was induced by 4-day exposure to Advanced DMEM/F12 (ThermoFisher #12634010) supplemented with L-glutamine (2 mM, ThermoFisher #25030081), Penicillin-Streptomycin (100 Units/ml; 100 μg/ml respectively, ThermoFisher #15140122), FGF4 (500 ng/ml, R&D Systems, #235-F4/CF), CHIR99021 (3 μM, Tocris, #4423) and defined FBS (2% vol/vol, GElifesciences, Hyclone #SH30070.02), which was refreshed daily. For the induction of human intestinal organoids, the adherent monolayer containing three-dimensional structures and the free-floating spheres that developed during mid/hindgut induction were mechanically harvested and fragmented by repeated pipetting. After gravity-based sedimentation of fragments, supernatant was discarded and the fragments were re-suspended in Basement Membrane Matrigel (Corning #354234) and plated in domes in a 24-well plate (ThermoFisher #142475). After 10 minutes of incubation at 37°C, Matrigel domes had solidified and were overlaid with Advanced DMEM/F12 (ThermoFisher #12634010) supplemented with L-glutamine (2 mM, ThermoFisher #25030081), Penicillin-Streptomycin (100 Units/ml; 100 μg/ml respectively, ThermoFisher # 15140122), Noggin (100 ng/ml, R&D Systems, #6057-NG/CF), EGF (100 ng/ml, R&D Systems, #236-EG), CHIR99021 (2 μM, Tocris, #4423), B27 (1x, ThermoFisher #17504044). Media was replaced every other day. After 7-10 days of culture, the density of the organoids was reduced by passaging: Matrigel domes were mechanically dislodged and organoids were released from the domes by repeated pipetting. The suspension was spun down at 1500 rpm for 3 minutes and the supernatant and Matrigel layer were aspirated. The organoids were washed twice in DPBS (Gibco, #14190-094), re-suspended in fresh Basement Membrane Matrigel and plated in domes as described before.

### Selection of epithelial cells from intestinal organoids

After 17-18 days of growing the human intestinal organoids in Matrigel domes, intestinal epithelial cells were selected to remove the majority of mesenchymal cells that co-develop with the differentiation. Matrigel domes were mechanically dislodged and organoids were released from the domes by repeated pipetting. The suspension was spun down at 1500 rpm for 3 minutes and the supernatant and Matrigel layer were aspirated, leaving only the organoid pellet. The organoids were washed twice in DPBS, re-suspended in warm TrypLE Select (ThermoFisher #12563029) and incubated for 5-6 minutes at 37 °C. The organoids were then dissociated by repeated pipetting. Depending on whether the organoids had sufficiently dissociated, the incubation with TrypLE Select was prolonged. To stop TrypLE Select activity after the organoids were dissociated to a single-cell suspension, DPBS supplemented with FCS (10% vol/vol, Gibco, #10270-106) was added to the cell suspension, which was thereafter passed through a 40-μm filter. The filtered cell suspension was spun down at 1500 rpm for 3 minutes and re-suspended in DPBS supplemented with FCS (10% vol/vol). Epithelial cells were enriched using the EasySep™ Human EpCAM Positive Selection Kit II (Stemcell Technologies, #17846) according to manufacturer’s instructions. The resulting epithelial cells were counted and cryopreserved in CryoStor CS10 (Stemcell Technologies, #7930) until seeding in the intestine-on-chip system.

### Seeding and maintenance of cells in the intestine-on-chip system

To develop the intestine-on-chip, we used the PDMS-based Chip-S1 and associated instrumentation from Emulate Inc (Boston, MA). This system contains two microfluidic channels, separated by a porous membrane. First, the PDMS surface inside the microfluidic channels was activated for extracellular matrix (ECM) coating according to the manufacturer’s instructions. Briefly, ER-1 solution (ER105, Emulate Inc) was introduced to the channels and the chips were exposed to UV light (36W, 365nm) for 10 minutes. This step was repeated once more with fresh ER-1 solution and 5-minute UV exposure. After UV treatment, the channels were washed with ER-2 solution (ER225, Emulate Inc) and subsequently with DPBS. After aspiration of DPBS, a solution of Basement Membrane Matrigel (83 µg/ml, Corning #354234) diluted in Advanced DMEM/F12 (ThermoFisher #12634010) was introduced to the channels and chips were stored at 4°C overnight. The next day (day −7), chips were incubated at room temperature for 1 hour before cell seeding. Intestinal epithelial cells were thawed, resuspended in Advanced DMEM/F12 (ThermoFisher #12634010) supplemented with L-glutamine (2 mM, ThermoFisher #25030081), Penicillin-Streptomycin (100 Units/ml; 100 μg/ml respectively, ThermoFisher # 15140122), Noggin (100 ng/ml, R&D Systems, #6057-NG/CF), EGF (100 ng/ml, R&D Systems, #236-EG), CHIR99021 (2 μM, Tocris, #4423), B27 (1x,

ThermoFisher #17504044), SB202190 (10 µM, Tocris #1264/10), A83-01 (500 nM, Tocris #2939/10), hereafter named ‘expansion medium’ (EM), and Y-27632 (10 µM, Tocris #1254/10). The chip channels were washed with EM supplemented with Y-27632 (10 µM) and cells were seeded at a concentration of 7.5 million/ml in the top channel. The remainder of cells that were not used for seeding chips, were fixed for use in flow cytometry analysis as described in section ‘Flow cytometry’. The chips were incubated for 2-3 hours in a humidified environment at 37 °C in 5% CO2 until cells were attached to the membrane. The top channel of the chip was gently washed with equilibrated EM supplemented with Y-27632 (10 µM) and chips were connected to the Emulate instrument and maintained in a humidified environment at 37 °C in 5% CO2. For the entire duration of the experiment, the flow rate of the media within both channels was 40 ul/hour, which imposes a wall shear stress on the cells of 0.067 mPa in the top channel and 1.7 mPa in the bottom channel. Culture media added to the chips was always first equilibrated. One day after seeding (day −6), fresh EM was supplied to the chips, which was repeated every 2-3 days. Media conditions were initiated 7 days after seeding of the chips (day 0): chips were either maintained in EM or differentiation medium (DM) was introduced to both channels or the top channel (EM-DM). DM was composed of Advanced DMEM/F12 (ThermoFisher #12634010) supplemented with L-glutamine (2 mM, ThermoFisher #25030081), Penicillin-Streptomycin (100 Units/ml; 100 μg/ml respectively, ThermoFisher #15140122), EGF (100 ng/ml, R&D Systems, #236-EG), B27 (1x, ThermoFisher #17504044), DAPT (10 µM, Tocris #2634) and PD0325901 (0.2 µM, Tocris #4192). Chips were maintained in these media conditions for 6 days (days 0-6) and respective media were refreshed every 2 days. Brightfield images were taken daily to assess tissue morphology in the chips over time.

### Barrier integrity analysis

One day after seeding the cells in the chip (day −6), 3-5 kDa fluorescein isothiocyanate– dextran (FITC-dextran; 500 µg/ml, Sigma-Aldrich #FD4-100MG) was added to the culture media in the top channel. Two hours after the introduction of FITC-dextran, the top and bottom channel outlet reservoirs of the chip system were emptied, corresponding to timepoint T=-6. For the duration of the experiment, every 24 hours all media was collected from the outlet reservoirs of the top and bottom channel and 50 µl culture media was collected from the inlet reservoirs of the top and bottom channel. The samples taken from the outlet reservoirs were weighed to determine the precise culture media volume that flowed through the top and bottom channel during the sampling interval and centrifuged (1000xg, 10 minutes, 4 °C) to remove cell debris. The fluorescence intensity in the supernatant samples, standard curve samples and fresh culture medium (blanc) was measured in black round-bottom 96-well plates at 485 nm excitation and 528 nm emission wavelength using the

BioTek Synergy HT plate reader. The standard curve included 14 dilutions (2000-0.24 µg/ml) of FITC-dextran in Advanced DMEM/F12. The raw fluorescence intensity values were converted to FITC-dextran concentration using the Emulate Standard Curve Calculator EC003v1.0^68^. The apparent permeability (Papp) was calculated using the formula: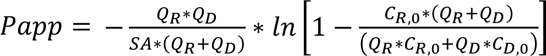, according to the Emulate Apparent Permeability Calculator EC004v1.0^69^, where Papp is in cm/s, the surface area of the co-culture channel (SA) is 0.171 cm^2^, Q_R_ and Q_D_ are the calculated fluid flow rates in the receiving (bottom) and dosing (top) channel, respectively, in cm^3^/s, C_R,0_ and C_D,0_ are the recovered concentrations in the receiving (bottom) and dosing (top) channel, respectively, in any consistent unit.

### Cytokine stimulation

On day 6, interferon (IFN)-γ (100 ng/ml, BioLegend #570204) or IFN-β (500 U/ml, PBL Assay Science #11410-2) was introduced to the culture media of the bottom channel of the intestine-on-chip in EM-DM condition, incubated for 6 hours and then subjected to dissociation for single-cell RNA sequencing.

### Immunofluorescent microscopy

On day 6, the tissue in the intestine-on-chip systems exposed to the EM, EM-DM and DM conditions was fixed to be used for immunofluorescent microscopy. The channels were washed twice with DPBS, paraformaldehyde (PFA, 4% vol/vol, ThermoFisher #28908) diluted in DPBS was introduced to the top and bottom channels and chips were incubated for 15 minutes at room temperature. The channels were washed three times with DPBS and chips were submerged in DPBS supplemented with Penicillin-Streptomycin (100 Units/ml; 100 μg/ml respectively, ThermoFisher #15140122) and stored at 4 °C until use for sectioning. To generate cross-sectional slices, thin sections of the PDMS were cut away on all four sides of the chip and chips were cut in half perpendicular to the channels using a microtome blade (pfm medical #207500002). Cross-sectional slices with a thickness of 200 µm were generated using a vibratome (VT1000S, Leica). During sectioning, chips were submerged in DPBS with ice and the following settings were used: speed=0.075 mm/s, frequency=60-65 Hz, blade angle: 5°, razorblades= Astra Superior Platinum Double Edge. The lower part of the blade holder of the vibratome was replaced by a manually designed piece with a flatter design, to prevent collision of this piece with the chip while sectioning. For immunofluorescent staining, the tissue slices were permeabilized by incubation in Triton X-100 (0.1% vol/vol, Sigma Aldrich #T8787) diluted in DPBS for 30 minutes at room temperature and blocked by incubation in bovine serum albumin (BSA, 3% weight/vol, Sigma Aldrich #A2153) diluted in DPBS for 1 hour at room temperature. Samples were stained overnight at 4 °C in BSA (3% weight/vol) diluted in DPBS containing the primary antibodies. The next day, samples were washed twice with DPBS and stained in BSA (3% weight/vol) diluted in DPBS containing the secondary antibodies for 2 hours at room temperature in the dark. Then samples were washed three times with DPBS and mounted on a glass microscope slide in a mounting medium with DAPI (Vector Laboratories #H-2000). Samples were kept at 4 °C in the dark until imaging. Images were taken using a Leica SP8 CLSM confocal immunofluorescent microscope (using the 10x or 20x objective). Results were analyzed using the Leica LAS X software. For every stained marker in each chip, slices were used from two different positions along the chip channel. Primary and secondary antibodies are listed in Table S3 and S4, respectively.

### Tissue height quantification

Brightfield images were taken of the cross-sectional slices used for immunofluorescent microscopy. The height of the tissue in the top channel of the intestine-on-chip was derived in Fiji-2 using this approach: For each chip slice, the largest and smallest distances from the tissue surface in the top channel straight to the chip membrane were measured, the average was calculated and the distance was converted to height in µm using the scale bar. For each biological replicate in each condition, the height of 8 cross-sectional slices, derived from different positions along the length of the chip channels, was measured using this method and the average was calculated. To derive the relative tissue height for the EM-DM and DM conditions, the percentage of the average tissue height relative to the height in EM (100%) was calculated.

### Dissociation of single-cells from the intestine-on-chip

On day 6, cells were dissociated from the intestine-on-chip systems exposed to the EM, EM-DM, DM, EM-DM + IFN-γ and EM-DM + IFN-β conditions to be used for single-cell RNA sequencing and flow cytometry. The top and bottom channels of the intestine-on-chip were washed with DPBS. TrypLE Select Enzyme solution (ThermoFisher #12563011) was introduced to the bottom channel and chips were incubated for 10 minutes at 37 °C, then triturated to detach the cells in the bottom channel. The cell suspension from the bottom channel of the chip was collected in a microcentrifuge tube, incubated for 5 minutes at 37 °C and further dissociated to a single cell suspension by gentle repeated pipetting. The single-cell suspension was added to a centrifuge tube containing Advanced DMEM/F12 (ThermoFisher #12634010) supplemented with FCS (10% vol/vol), Y-27632 (10 µM) and Deoxyribonuclease II (10 U/ml, Sigma Aldrich #D8764-30KU), hereafter named ‘dissociation solution’, and stored on ice. The bottom channel of the chip was washed with DPBS and TrypLE Select Enzyme solution was introduced to the top channel. Chips were incubated for 10 minutes at 37 °C, then triturated to detach the cells in the top channel. Detached tissue was collected in a microcentrifuge tube, incubated for 5 minutes at 37 °C and further dissociated to a single cell suspension by gentle repeated pipetting. The single-cell suspension was added to a centrifuge tube containing ‘dissociation solution’ and stored on ice. Fresh TrypLE Select Enzyme solution was added to the top channel of the chip and the previously described steps were repeated (3-4 times) until the tissue was removed from the top channel of the chip. The cell suspensions of the top and bottom channel were combined and passed through a 70-μm filter. The filtered suspension was centrifuged (300xg, 5min, 4 °C) and resuspended in ‘dissociation solution’. The cells were counted using a hemocytometer and viability was assessed using Trypan Blue (Sigma Aldrich #T8154) staining.

### Flow cytometry

From each intestine-on-chip, 0.5 million dissociated cells were used in flow cytometry analysis. A fraction of the cell suspension of the biological replicates in each condition was pooled together to be used as ‘unstained sample’ and ‘single-antibody stained samples’ in flow cytometry analysis. The remainder of the cells were labelled using the Zombie Aqua Fixable Viability Kit (BioLegend #423101) according to the manufacturer’s instructions to indicate the viability status of each cell. Subsequently, the unstained and Zombie Aqua-stained samples were fixed by incubation in PFA (4% vol/vol, ThermoFisher #28908) diluted in DPBS for 10 minutes at room temperature, washed with DPBS and stored in DPBS with FCS (2% vol/vol) at 4 °C until flow cytometry analysis. The same procedure was performed with the cell suspension samples which were taken during seeding of the intestine-on-chip. The cells were permeabilized and labelled with fluorophore-conjugated antibodies using the BD Perm/Wash buffer (BD Biosciences # 554723) according to the manufacturer’s instructions. BD Horizon Brilliant Stain Buffer (BD Biosciences #563794) was included in the antibody mixture in an equal volume to the total volume of the antibodies. The cells were resuspended in DPBS with FCS (2% vol/vol) and analyzed using the Cytek Aurora. Results were analyzed using the Kaluza Analysis software (Beckman Coulter Life Sciences). Unstained and ‘fluorescence-minus-one’ samples were included in the analysis as the basis for the gating strategy (Figure S3B). Single-antibody stained samples were used for compensation. Antibodies are listed in Table S5.

### Single-cell library preparation and sequencing

Five pools were prepared from the dissociated cell suspensions with a final concentration of 1000 cells/µl per pool: Each pool consisted of the three biological replicates, to later segregate based on genotype, and the conditions were balanced over the pools. Single cells were captured with the 10X Chromium controller (10x Genomics) according to the manufacturer’s instructions (document CG000315 Rev B) and as described earlier^70^. Each sample pool was loaded into a different lane of the same 10X chip (Chromium Next GEM Chip G Single Cell Kit, PN-1000127, 10x Genomics). The cDNA libraries were prepared using the Chromium Next GEM Single Cell 3’ Kit v3.1 (Dual Index) (PN-1000269, 10x Genomics) reagents according to the manufacturer’s instructions (document CG000315 Rev B). The libraries were sequenced with a targeted sequencing depth of 50.000 reads per cell on an MGISEQ-2000 using a 150-bp paired-end kit and following the 10x Genomics guidelines (BGI Tech Solutions, Hong Kong).

### Single-cell RNA sequencing data processing and analysis

#### Processing of sequencing data and cell-type annotation

CellRanger 6.1.2 software with default settings was used to demultiplex the sequencing data, generate FASTQ files, align the sequencing reads to the human reference genome (GRCh38), filter cell and unique molecular identifier (UMI) barcodes, and count gene expression per cell. Doublet detection was done using SoupOrCell v2^71^ making use of the 1000g common variants to confine the search space. The genotypic clusters in each 10X lane identified by Souporcell were subsequently correlated to the GSA genotypes of the three donors and assigned to donors based on the best matching correlation. RStudio with the R package Seurat v4.1.1 was used for data preprocessing and downstream analysis^72^. Cells with a high proportion of mitochondrial genes (≥20%), and the cluster of cells that these cells were primarily assigned to, were discarded, as this is an indicator of low-quality cells^73^. Additionally, cells expressing ≤ 2000 genes or ≥ 100.000 UMI counts per cell were considered outliers and discarded. Finally, all cells that were marked as doublets or inconclusive by SoupOrCell v2^71^ (marking inter-sample doublets) or cells associated with these doublets based on similarity in gene expression by scDblFinder^74^ (marking intra-sample doublets) were discarded. After filtering, the datasets contained 3275 (EM), 4170 (EM-DM) and 3658 (DM), 3955 (EM-DM + IFN-β) and 3050 (EM-DM + IFN-γ) cells and were used for downstream analysis. The datasets of the biological replicates were aligned and integrated using Seurat’s reciprocal PCA method. The dataset was normalized using Seurat’s SCTransform (vst.flavor = ‘v2’) function and dimension reduction was performed using principal component analysis (PCA) and uniform manifold approximation and projection (UMAP) on the first 30 principal components (PCs). Then, cell clustering was performed using Seurat’s FindNeighbors (dims = 1:30) and FindClusters function (resolution 0.34). The resulting clusters were visualized using a UMAP plot and annotated to well-established intestinal cell types by the expression of canonical cell-type markers. Each cell-type cluster was assigned to the epithelial or mesenchymal/neural compartment based on the average expression of compartment-characteristic genes *EPCAM* and *VIM (*Epithelial cell: *EPCAM* ≥ 1.0 and *VIM* < 2.7; mesenchymal/neural cell: *EPCAM* < 1.0 and *VIM* ≥ 2.7). Seurat’s CellCycleScoring function was used to project the cell cycle phase of each cell. Cells in the ‘S’ and ‘G2M’ phase were designated ‘Dividing’ and cells in the ‘G1’ phase were designated ‘Non-dividing’. After cell type, compartment and cell cycle phase annotation, two subsets of the dataset were generated: (1) the data of the intestine-on-chip exposed to EM, EM-DM and DM, (2) the data of the intestine-on-chip exposed to EM-DM, EM-DM + IFN-γ and EM-DM + IFN-β. The first dataset was used for Figures 3-5 and S4-7 and Tables S1 and S2 and the second dataset was used for Figures 6 and S8.

#### Differential gene expression and pathway analysis

For differential gene expression (DE) analysis we used Seurat’s FindMarkers function on the corrected counts stored in the data slot of the SCTransform-normalized data (test.use = “MAST”, min.pct = 0.1, logfc.threshold = 0.25). The variables ‘donor’ and ‘cellular detection rate’^75^ were included as variables to test (latent.vars) in the FindMarkers function, after comparing the effect of potential confounding variables on the p-value distribution of DE genes using the MAST package^75^. To compare cell types, one cell type was compared to the rest of the data. To compare media conditions, pairwise DE analysis was performed, providing three lists of DE genes: EMvsEM-DM, EMvsDM, EM-DMvsDM. To compare IFN stimulation conditions, two pairwise DE analyses were performed comparing EM-DM + IFN-γ or EM-DM+ IFN-β to EM-DM. The relevant cellular compartments were selected for each analysis as indicated in the figures. Volcano plot visualizations of the −log10(adjusted p-value) and average log2 fold change (log2FC) were used to determine thresholds to filter significant DE genes: DE genes with adjusted p-values < 0.01 (considered statistically significant) and an average log2FC > 0.5 were selected for downstream analysis. The absolute value of the average log2FC was used to filter DE genes when comparing media conditions to include genes upregulated in all conditions. The average expression levels of DE genes in specific groups of the data were visualized using Seurat’s DotPlot function or using the Heatmap function of the package ComplexHeatmap after performing hierarchical clustering to cluster the genes. The overlap of DE genes between conditions was visualized using the venn.diagram function of package VennDiagram. The DE genes were used to identify pathways, hereby distinguishing up- and downregulated genes if relevant, using the clusterProfiler v4.6.2 package^76^ and the ‘Gene Ontology: biological processes’^77,78^ or ‘Reactome’^79^ database, as indicated in the figure caption. The five pathways with the lowest adjusted p-value were selected per group and the detected gene ratio and adjusted p-values were projected using the dotplot function from the enrichplot package. For the pathway enrichment analysis of the cell-type clusters, an average log2FC > 1.0 was used as threshold to filter DE genes to display the most characteristic pathways per cell subtype (Figure S4C).

#### Cell-type abundance

The proportion of cell subtypes within groups was analyzed using the propeller function (transform=’logit’) of the Speckle package^80^. Using the same function, compositional differences between groups were tested on statistical significance: ANOVA was instigated within the propeller function, as more than 2 groups were tested. The results were visualized as barplot.

#### Comparison to reference data

To compare the intestine-on-chip dataset to the human intestine, we mapped our dataset to the Gut Cell Atlas dataset^27,31^, containing 428.469 intestinal cells from different age groups (first and second trimester (fetal), pediatric and adult), intestinal regions (appendix, rectum, mesenteric lymph node, small intestine and large intestine) and cell types (epithelial, mesenchymal, neural, endothelial and immune cells). We used Seurat’s reference mapping method, specifically the FindTransferAnchors and MapQuery functions, to verify our cell-type annotation and predict the age group, category, and intestinal region of each cell in our dataset using the following projected variables: ‘Region’, ‘cell_type’, ‘category’, ‘Age_group’. The proportions of cell subtypes were generated separately for the intestine-on-chip and Gut Cell Atlas datasets using the propeller function of the Speckle package as described before and visualized in one barplot. For the Gut Cell Atlas, the individual samples, of which the average was calculated per group, were defined as the combination of the donor code (variable ‘Sample.name’) and the region code (variable ‘Region.code’). For the cell-type composition comparisons between the intestine-on-chip and Gut Cell Atlas dataset, the original annotations of each dataset were used. To assess the similarity between the intestine-on-chip dataset and the Gut Cell Atlas dataset, correlation scores were generated between groups in each dataset. First, the intestine-on-chip and Gut Cell Atlas datasets were normalized and log1p-transformed using Seurat’s NormalizeData function. Then, the normalized data was used to generate a matrix of the average expression of the selected DE genes (as indicated in the figure caption) per group. Gene expression was then mean-centered to 0 and scaled per-gene to standard deviation of 1. The resulting expression matrices of the intestine-on-chip and Gut Cell Atlas datasets were combined and Pearson correlation scores and p-values were generated using the rcorr function of the Hmisc package. These scores were visualized using the corrplot function of the corrplot package, in which color intensity and dot size corresponded to correlation score ‘r’ and non-significant correlations (p-value > 0.01) were left blank.

For the analyses with the Gut Cell Atlas reference dataset, subsets of the data were used as follows: The inflammatory bowel disease-related samples (Diagnosis = ‘Pediatric Crohn Disease’) were excluded in all analyses. The resulting dataset including all age groups, regions and cell types was used for the reference mapping displayed in Figures 4A-C, 4E and S6B. For all further analysis (correlation and cell-type composition), a subset of the data was used corresponding to small intestine and epithelial, mesenchymal and neural cells as indicated in the figures (Region = ‘SmallInt’ and category = ‘Epithelial’, ‘Mesenchymal’, ‘Neuronal’) (Figures 4D and 4F-I). Different age groups were used per analysis, as indicated in the figures. For the composition analysis in Figure 4F and correlation analysis in Figure 4I, the samples of the Gut Cell Atlas data that were subjected to an enrichment of *EPCAM*-positive cells (some fetal first trimester samples and all pediatric samples) were not included, to prevent proportion values or correlation scores, respectively, to be the result of the altered ratio of EPCAM-positive and negative fractions. In these analyses, the resulting first and second trimester samples that were not sorted based on *EPCAM* expression were combined as ‘fetal’. For the composition analysis in Figures 4F-H, one adult donor (A32 (411C)) was excluded based on the aberrant Paneth cell ratio of 50% from all epithelial cells in an ileal sample.

The cell-type annotations of the Gut Cell Atlas dataset were simplified or detailed according to the description in Elmentaite et al.^27^ as follows: ‘Enteroendocrine cell’ was originally ‘D cells (SST+)’, ‘EC cells (NPW+)’, ‘EC cells (TAC1+)’, ‘EECs’, ‘I cells (CCK+)’, ‘K cells (GIP+)’, ‘L cells (PYY+)’, ‘M/X cells (MLN/GHRL+)’, ‘N cells (NTS+)’, ‘β cells (INS+)’ or ‘Progenitor (NEUROG3+)’ (original annotation used in Figure 4E); ‘Fetal proximal progenitor’ was originally ‘Proximal progenitor’; ‘Fetal distal progenitor’ was originally ‘Distal progenitor’; ‘Fetal CLDN10+ cells’ was originally ‘CLDN10+ cells’; ‘Goblet cell’ was originally ‘Goblet cell’ or ‘BEST2+ Goblet cell’; ‘Neuron’ was originally ‘Branch A1 (iMN)’, ‘Branch A2 (IPAN/IN)’, ‘Branch A3 (IPAN/IN)’, ‘Branch A4 (IN)’, ‘Branch B1 (eMN)’, ‘Branch B2 (eMN)’ or ‘Branch B3 (IPAN)’; ‘Glial cell’ was originally ‘Glia 1 (DHH+)’, ‘Glia 2 (ELN+)’, ‘Glia 3 (BCAN+)’, ‘Differentiating glia’ or ‘Adult Glia’; ‘Stromal subtype’ was originally ‘Stromal 1 (ADAMDEC1+)’, ‘Stromal 1 (CCL11+)’, ‘Stromal 2 (CH25H+)’, ‘Stromal 2 (NPY+)’, ‘Stromal 3 (C7+)’, ‘Stromal 3 (KCNN3+)’, ‘Transitional Stromal 3 (C3+)’ or ‘Stromal 4 (MMP1+)’; ‘Interstitial cell of Cajal’ was originally ‘ICC’; ‘Smooth muscle cell’ was originally ‘SMC (PLPP2+)’ or ‘SMC (PART1/CAPN3+)’; ‘Mesoderm’ was originally ‘Mesoderm 1 (HAND1+)’ or ‘Mesoderm 2 (ZEB2+)’; ‘Mesothelium’ was originally ‘Mesothelium’, ‘Mesothelium (PRG4+)’ or ‘Mesothelium (RGS5+)’; ‘Lymph node fibroblast’ was originally ‘mLN Stroma (FMO2+)’, ‘T reticular’, ‘mLTo’ or ‘FDC’; ‘Myofibroblast’ was originally ‘myofibroblast’ or ‘myofibroblast (RSPO2+)’; ‘Pericyte’ was originally ‘angiogenic pericyte’, ‘Contractile pericyte (PLN+)’, ‘Immature pericyte’ or ‘Pericyte’.

The intestine-on-chip data was compared to the Human Fetal Endoderm Atlas to verify the intestinal phenotype of our dataset, using the score_fidelity function in the scoreHIO package^30^.

#### Trajectory analysis

Trajectories between cell types were determined based on RNA velocity analysis and transcription similarity estimation (Python-based using package scanpy 1.9.3^81^). First, velocyto was used to calculate spliced and unspliced RNA counts^82^. The pre-processed Seurat object containing normalized counts (SCT, slot data) was converted to AnnData and merged with the spliced/unspliced counts matrixes. RNA velocity was then calculated using scVelo 0.3.0, following their recommended workflow^83^. In short, the dataset was processed using the scv.pp.filter_and_normalize (min_shared_counts=20, n_top_genes=2000) and scv.pp.moments (n_pcs=30) functions. Velocities were calculated using the dynamical model, while accounting for competing kinetic regimes between cell types by running the following functions: scv.tl.recover_dynamics, scv.tl.velocity (mode = ‘dynamical’) and scv.tl.velocity (diff_kinetics=True). The velocities were visualized using the scv.pl.velocity_embedding_stream function and latent time was recovered using the scv.tl.latent_time function. The RNA velocity data was combined with a Partition-based graph abstraction (PAGA) analysis to identify and visualize trajectories using the scv.tl.paga and scv.pl.paga functions respectively.

To identify putative driver genes for each lineage in the intestine-on-chip, we used CellRank 2.0.0 with a combined VelocityKernel (weight of 0.8) and ConnectivityKernel (weight of 0.2)^84^. In short, eight macrostates were calculated by computing a Schur decomposition (n_components=20, method=’brandts’) using the Generalized Perron Cluster Cluster Analysis (GPCCA) estimator^85^. Seven of these macrostates were set as terminal state: “Enterocyte type 1”, “Enterocyte type 2”, “Paneth-like cell”, “Enteroendocrine cell”, “Mesenchymal-like epithelial cell type 2”, “Myofibroblast” and “Neuron”. Then, cellular fate was investigated by inferring fate probabilities for each predicted terminal state using the g.compute_fate_probabilities function. Next, fate probabilities and gene expression levels were correlated to identify putative driver genes for each terminal state, considering the relevant cell types in each lineage, using the g.compute_lineage_drivers function. The temporal activation of the top 40 putative driver genes along the pseudotime axis was visualized for each lineage by using the cr.pl.heatmap function and the following parameters: a model for GAM fitting (cr.models.GAM), MAGIC-imputed data^86^ and the latent time (latent_time) generated using the scVelo package as described before. To visualize the driver genes for each lineage, the relevant subset of the data (epithelial or mesenchymal/neural) was used.

#### Cell-cell communication analysis

Potential cell-cell interactions between the cell types in the intestine-on-chip were predicted based on the expression of ligands and receptors, taking into account the subunit organization of heteromeric complexes, using the CellPhoneDB v2.0 method^87^ and database implemented in the Liana package v0.1.12^88^. The following parameters were inherent in the method: an interaction was considered if the ligand and receptor genes were expressed in at least 10% of the cells, for heteromeric complexes the subunit with the minimum expression was used. The predicted interactions were filtered based on significance (p-value ≤ 0.01). Pathways were assigned to the resulting ligand-receptor combinations using the CellChatDB human database^89^. The contribution of each pathway to the predicted interactions in the intestine-on-chip was visualized as proportion of the total predicted interactions per media condition. The pathways with a contribution lower than 2% in all media conditions were named ‘other’ and closely related pathway families were combined: ‘EPHA’ and ‘EPHB’ were named ‘EPH’, ‘SEMA3’ and SEMA4’ were named ‘SEMA’, ‘ncWNT’ and ‘WNT’ were named ‘WNT’. For each pathway, potential sender and receiver cell types were visualized by plotting the frequency of receptor-ligand interactions between each pair of cell subtypes using Liana’s heat_freq function.

#### Cell-type annotation of interferon-responsive cells

IFN-β- and IFN-γ-responding cells were annotated based on the expression of stimulation-specific DE genes as determined based on DE analysis. IFN-β-responding cells were cells in the EM-DM + IFN-β dataset which complied with the following gene expression level thresholds: IFIT1 > 1, MX1 > 1.5 and IFI6 > 1. IFN-γ-responding cells were cells in the EM-DM + IFN-γ dataset which complied with the following gene expression level thresholds: IRF1 > 2, APOL6 > 1 and TAP1 > 1. Cells which were not annotated as IFN-β- or IFN-γ-responding cells were named IFN-β- or IFN-γ non-responding cells in the respective dataset. To identify the cell-type identity of the IFN-responding and non-responding cells, the cell types were re-annotated after correction for gene-expression differences resulting from the stimulation condition. To this extent, the datasets of each stimulation condition (EM-DM, EM-DM + IFN-β and EM-DM + IFN-γ) and biological replicate were aligned and integrated using Seurat’s reciprocal PCA method. The dataset was normalized using Seurat’s SCTransform (vst.flavor = ‘v2’) function and dimension reduction was performed using principal component analysis (PCA) and uniform manifold approximation and projection (UMAP) on the first 30 principal components (PCs). Then, cell clustering was performed using Seurat’s FindNeighbors (dims = 1:30) and FindClusters function (resolution 0.4). Each cluster was assigned the cell type that was the most prevalent according to the initial cell-type annotation.

### Statistics

The data was presented as mean± standard deviation or median with interquartile range. Significant differences between the three media conditions were determined using a one-way analysis of variance (ANOVA) test with the Tukey multiple comparisons test. Differences between groups were considered statistically significant when P-value <0.05. Rstudio with the R package rstatix was used for statistical analysis^90^.

## Data and code availability

The single-cell RNA-sequencing data and code or information required to reproduce the analysis detailed above will be shared by the lead contact upon request after publication.

## Acknowledgements

This work was supported by the Netherlands Organ-on-Chip Initiative, an NWO Gravitation project (024.003.001) funded by the Ministry of Education, Culture, and Science of the government of the Netherlands, an NWO Spinoza Prize (NWO SPI 92-266) and the United European Gastroenterology Research Prize to C.W. J.M is supported by a PhD scholarship from the Graduate School of Medical Sciences, University of Groningen. L.F. is supported by a grant from the Dutch Research Council (grant no. ZonMW-VICI 09150182010019), an ERC Starting Grant (grant agreement 637640 (ImmRisk)), an Oncode Senior Investigator grant, sponsored research collaborations with Biogen, Roche and Takeda and also received support from the European Union’s Horizon Europe Research and Innovation Programme under grant agreement No 101057553. I.H.J. is supported by a Rosalind Franklin Fellowship from the University of Groningen and an NWO VIDI grant (016.171.047). We thank Kate Mc Intyre for editing the manuscript. We thank the following facilities of the University Medical Center Groningen for their support and services: iPSC/CRISPR facility, UMCG Microscopy and Imaging Center and Flow Cytometry Unit. We thank Emulate Inc (Boston, USA) for kindly providing the Human Emulation System.

## Author contributions

R.M. designed experiments, analyzed experimental and single-cell RNA-sequencing data and wrote the manuscript. R.M., J.M. and A.D. developed experimental set-up and executed experiments. R.O. processed raw single-cell RNA-sequencing data. L.F. supervised R.O. C.W. assisted in conceiving the project. R.B. provided support on the experimental set-up. I.J. and S.W. supervised the project and provided feedback on experiment design, data analysis and writing.

## Supplemental information

**Supplementary Figure 1** - The development of villus-like folds and quantitative measures of the intestine-on-chip.

**Supplementary Figure 2** - Morphological differences induced by the EM, EM-DM and DM conditions.

**Supplementary Figure 3** - Microscopy and flow cytometry data of the cell-type composition in the intestine-on-chip.

**Supplementary Figure 4** - The characterization of cell-type clusters identified in the single-cell RNA sequencing data.

**Supplementary Figure 5** - Cell-type morphology, variability among donors and cell cycle states.

**Supplementary Figure 6 -** Comparison of the intestine-on-chip with the human intestine.

**Supplementary Figure 7** - Differentiation trajectories and cell-cell communication analysis in the intestine-on-chip.

**Supplementary Figure 8** - Characterization of the cellular response to IFN-β and IFN-γ in the intestine-on-chip.

**Supplementary Table 1** - Cell subtype abundance in the different media conditions in the intestine-on-chip.

**Supplementary Table 2** - Epithelial cell subtype proportions in the intestine-on-chip compared to the human adult small intestine.

**Supplementary Table 3** - Primary antibodies used for immunofluorescent microscopy.

**Supplementary Table 4** - Secondary antibodies used for immunofluorescent microscopy.

**Supplementary Table 5** - Antibodies used for flow cytometry.

