## Supplemental information for "An iPSC-derived small intestine-on-chip with self-organizing epithelial, mesenchymal and neural cells"

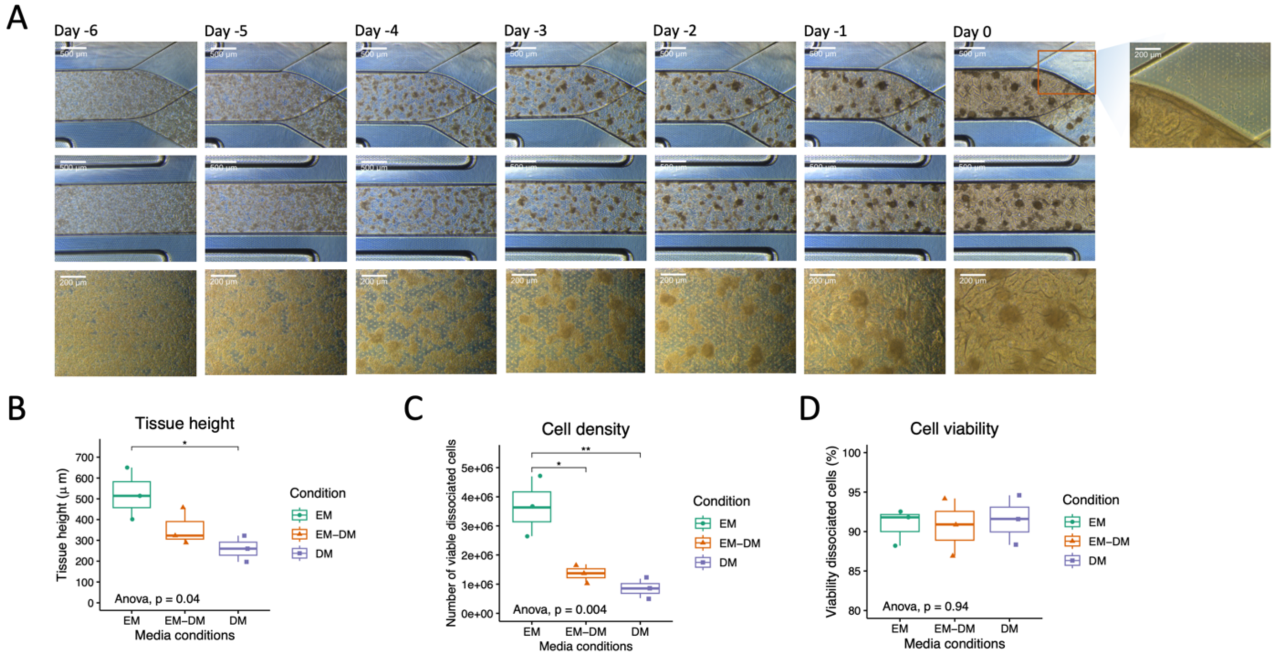


**Supplementary Figure 1. The development of villus-like folds and quantitative measures of the intestine-on-chip.** (A) Representative brightfield images of the top view of the intestine-on-chip during the first seven days after seeding (Days -6 to 0). The presence of cells in the bottom channel at day 0 is highlighted. (B) Quantification of tissue height of the tissue in the top channel at day 6, displayed as median values from three biological replicates. (C, D) The cell density and viability after dissociation of the cells from the top and bottom channels at day 6, displayed as median values from three biological replicates. P-value ≤ 0.05 (*), 0.01 (**), 0.001 (***), 0.0001 (****).

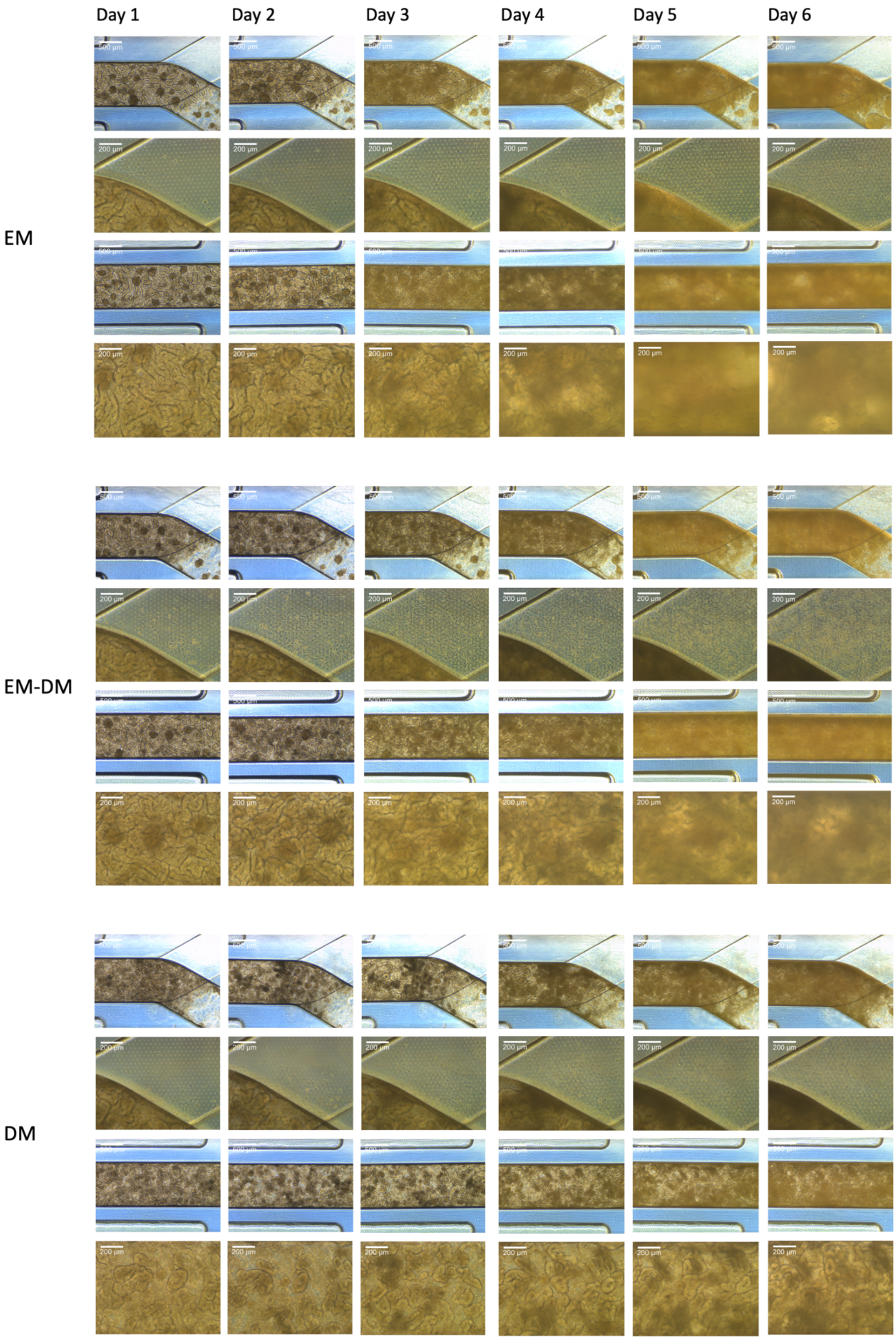


**Supplementary Figure 2. Morphological differences induced by the EM, EM-DM and DM conditions.** Representative brightfield images of the top view of the intestine-on-chip after introduction of the EM, EM-DM and DM media conditions (Days 1 to 6).

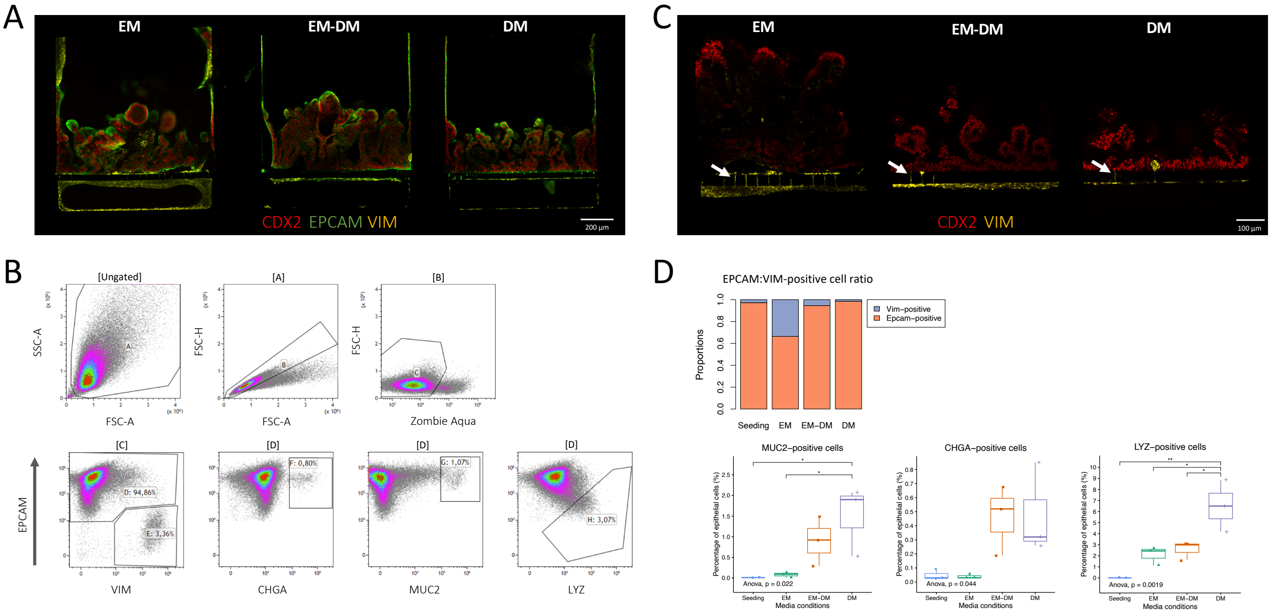


**Supplementary Figure 3. Microscopy and flow cytometry data of the cell-type composition in the intestine-on-chip.** (A) Representative immunofluorescent confocal images of cross-sectional slices of the intestine-on-chip stained for markers characteristic of intestinal epithelial cells (*EPCAM* and *CDX2*) and mesenchymal cells (*VIM*). (B) Gating strategy used in the analysis of the flow cytometry data. Cells were gated to discard cellular debris [A] and select single cells [B], viable cells [C], *EPCAM*-positive cells [D], *VIM*-positive cells [E], *CHGA*-positive cells [F], *MUC2*-positive cells [G] and *LYZ*-positive cells [H]. (C) Representative immunofluorescent confocal images of cross-sectional slices of the intestine-on-chip stained for markers characteristic of intestinal epithelial cells (*CDX2*) and mesenchymal cells (*VIM*). *VIM*-positive cells within the pores of the membrane separating the top and bottom channel are marked by arrows. (D) Quantification of cell-type proportions based on flow cytometry analysis, displayed as median values of three biological replicates. P-value ≤ 0.05 (*), 0.01 (**), 0.001 (***), 0.0001 (****).

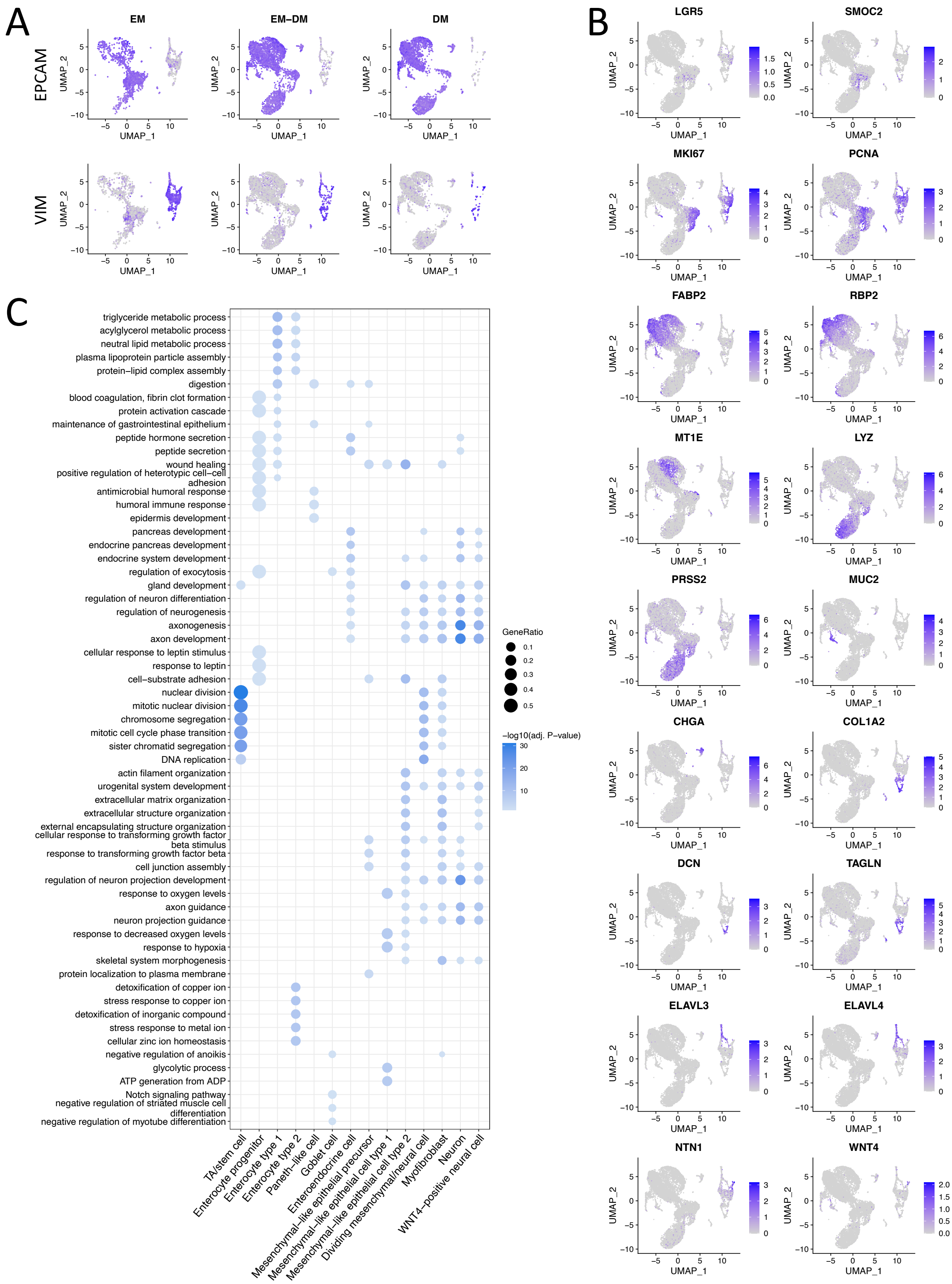


**Supplementary Figure 4. The characterization of cell-type clusters identified in the single-cell RNA sequencing data.** (A) UMAP projection of the expression of epithelial (*EPCAM*) and mesenchymal/neural (*VIM*) marker genes. (B) UMAP projection of the expression of canonical marker genes for intestinal epithelial, mesenchymal and neural subtypes: stem cells (*LGR5, SMOC2*), dividing cells (*MKI67, PCNA*), enterocytes (*FABP2, RBP2, MT1E*), Paneth cells (*LYZ, PRSS2*), goblet cells (*MUC2*), enteroendocrine cells

(*CHGA*), myofibroblasts (*COL1A2, DCN, TAGLN*), neural cells (*ELAVL3, ELAVL4, NTN1*) and *WNT4*. (C) Pathway enrichment of the differentially expressed genes between cell types using the Gene Ontology: biological process database.

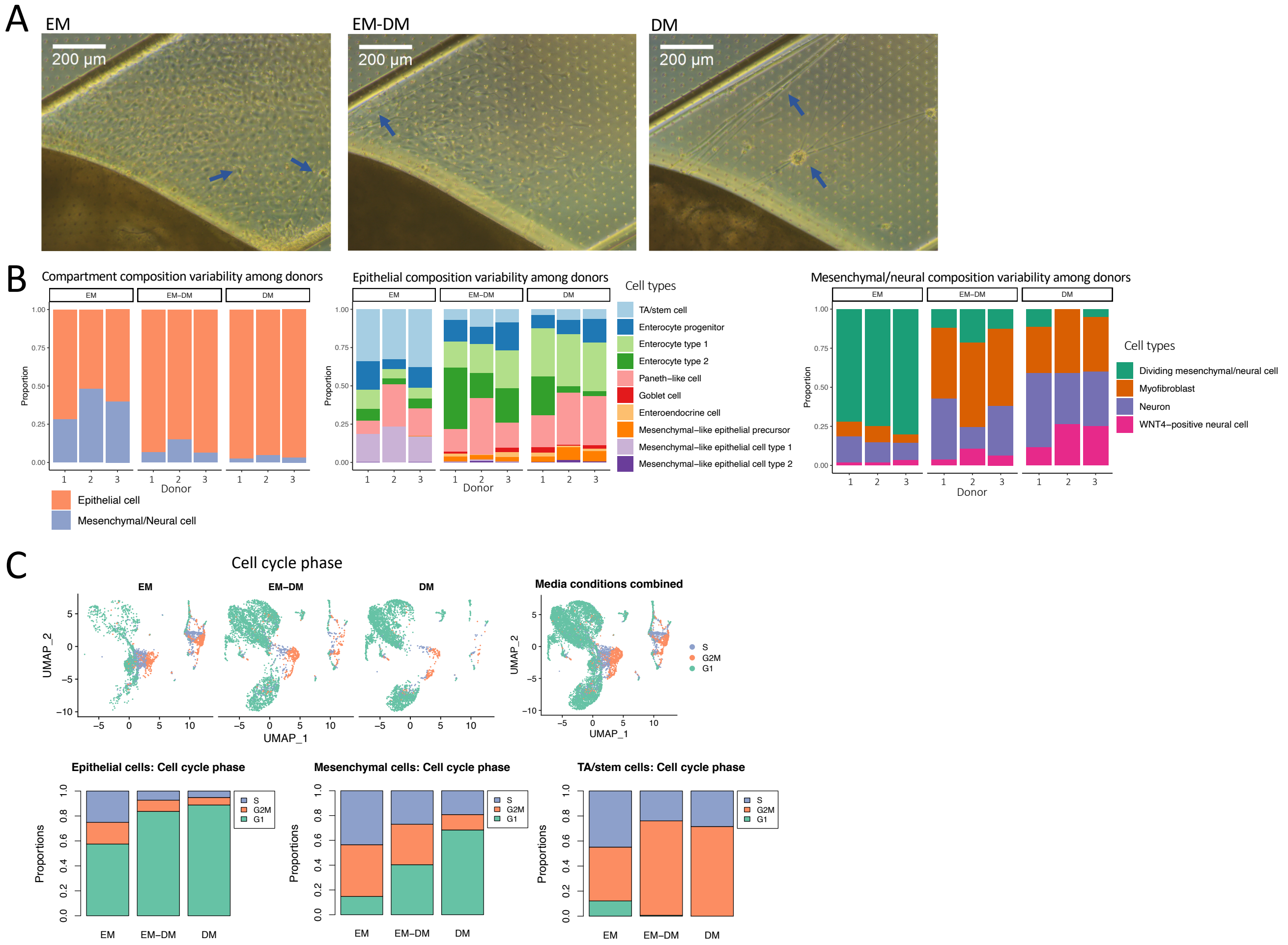


**Supplementary Figure 5. Cell-type morphology, variability among donors and cell cycle states.** (A) Brightfield images of the top view of the bottom channel in the intestine-on-chip at day 6. Cells in the bottom channel that have a morphology characteristic of neurons are marked by arrows. (B) Cell-type composition variability among the three donors. (C) UMAP projection and proportions of the cell cycle state of the cells in the intestine-on-chip.

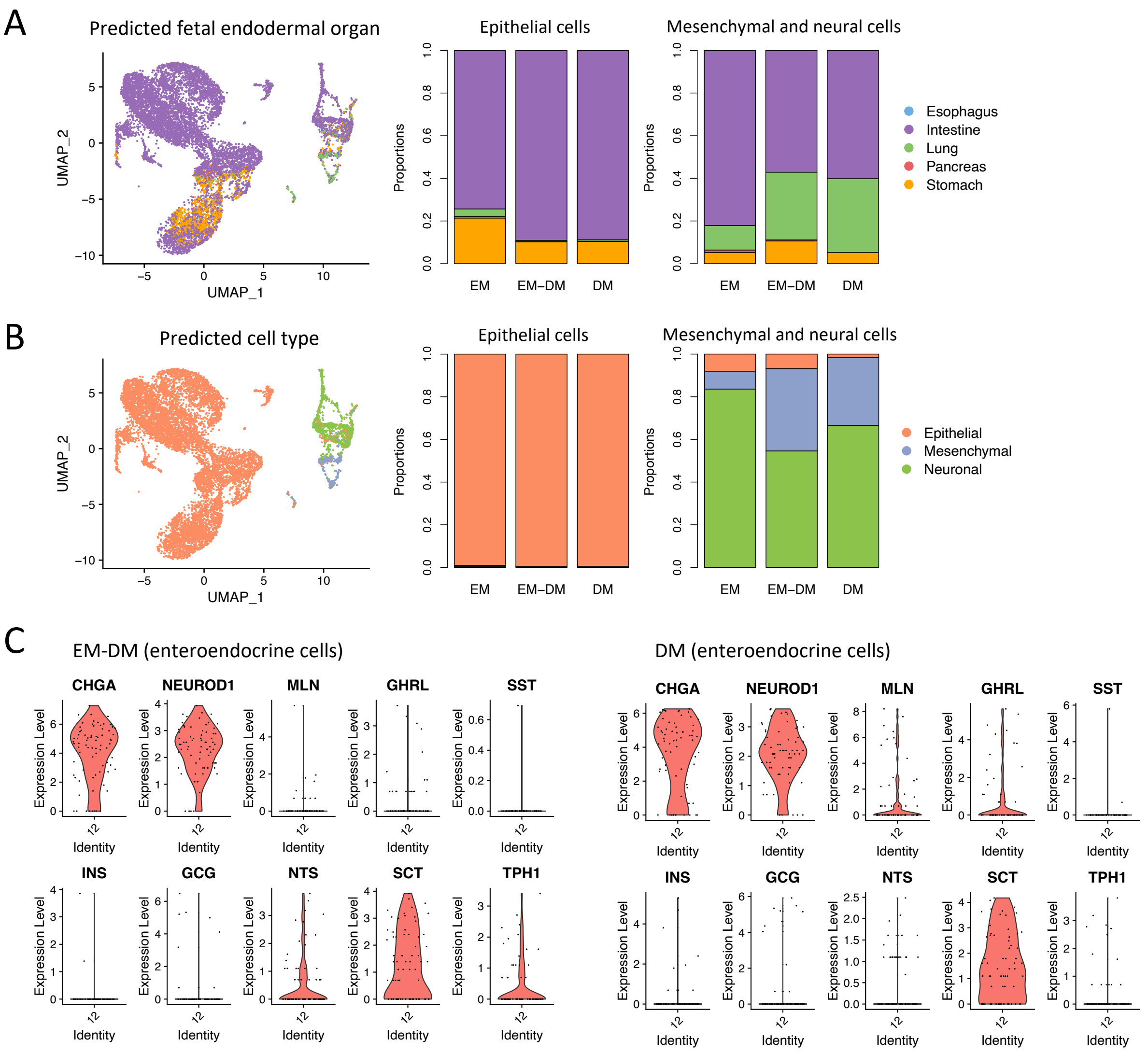


**Supplementary Figure 6. Comparison of the intestine-on-chip with the human intestine.** (A) UMAP projection and proportions of the predicted fetal endodermal organ based on the Human Fetal Endoderm Atlas. (B) UMAP projection and proportions of predicted cellular compartment based on human intestinal data of the Gut Cell Atlas (including the age groups first trimester, second trimester, pediatric and adult; the regions small intestine, large intestine, appendix, rectum and mesenteric lymph node; and the compartments epithelial, mesenchymal, neural, endothelial and immune cells). (C) Gene expression levels of intestinal hormones in the enteroendocrine cell cluster exposed to the EM-DM and DM conditions.

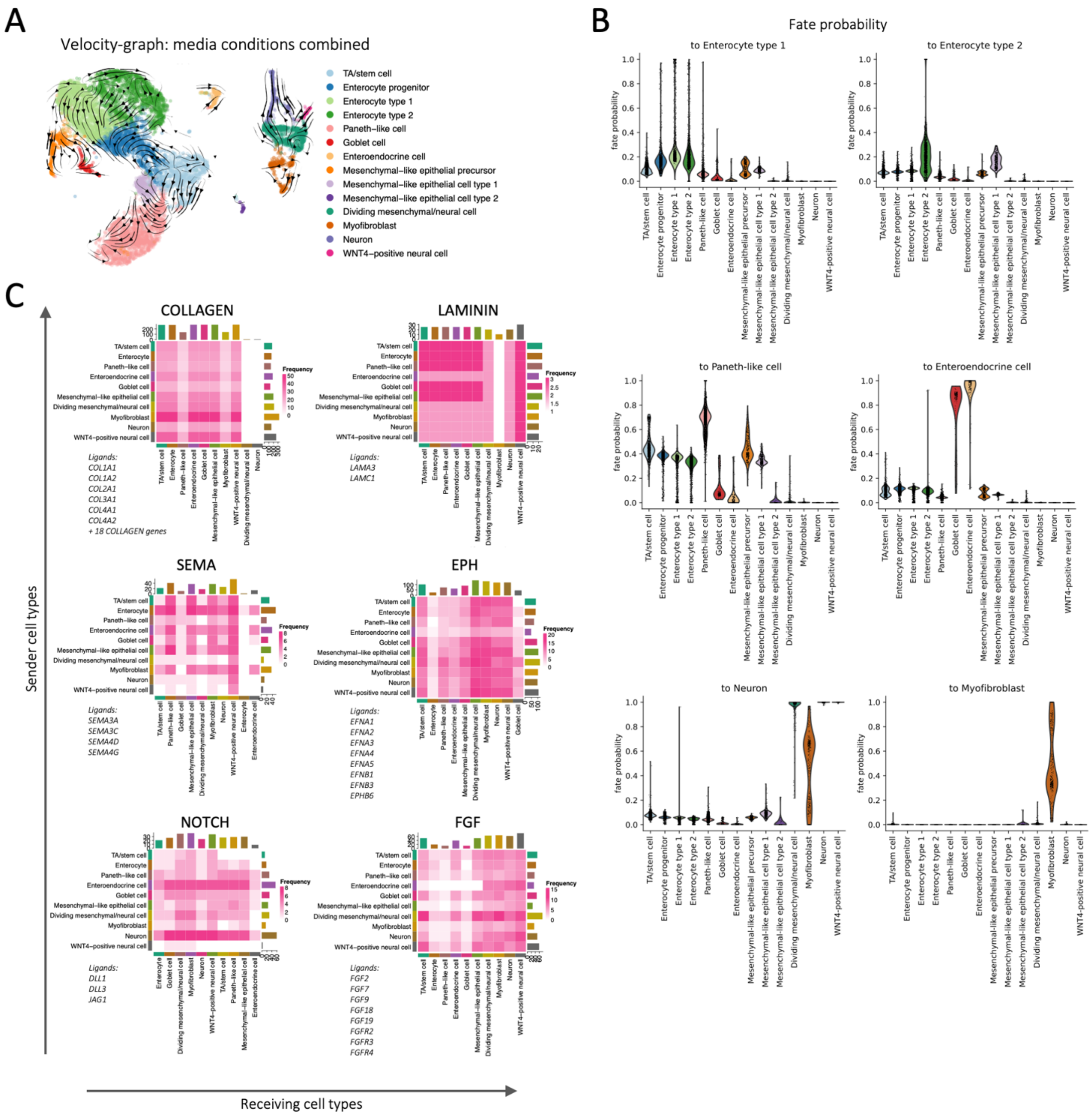


**Supplementary Figure 7. Differentiation trajectories and cell-cell communication analysis in the intestine-on-chip.** (A) Differentiation trajectories of cell types in the intestine-on-chip projected by RNA velocity graphs with overlaid arrows (scVelo). (B) Fate probability to contribute to a certain lineage, displayed using a Violin plot (scVelo and CellRank). (C) Frequency of potential communication events between cell types for the pathways with the most detected interactions determined by ligand-receptor expression analysis (Liana, CellPhoneDB). The analysis was performed using the combined dataset of the media conditions. The detected ligands are shown.

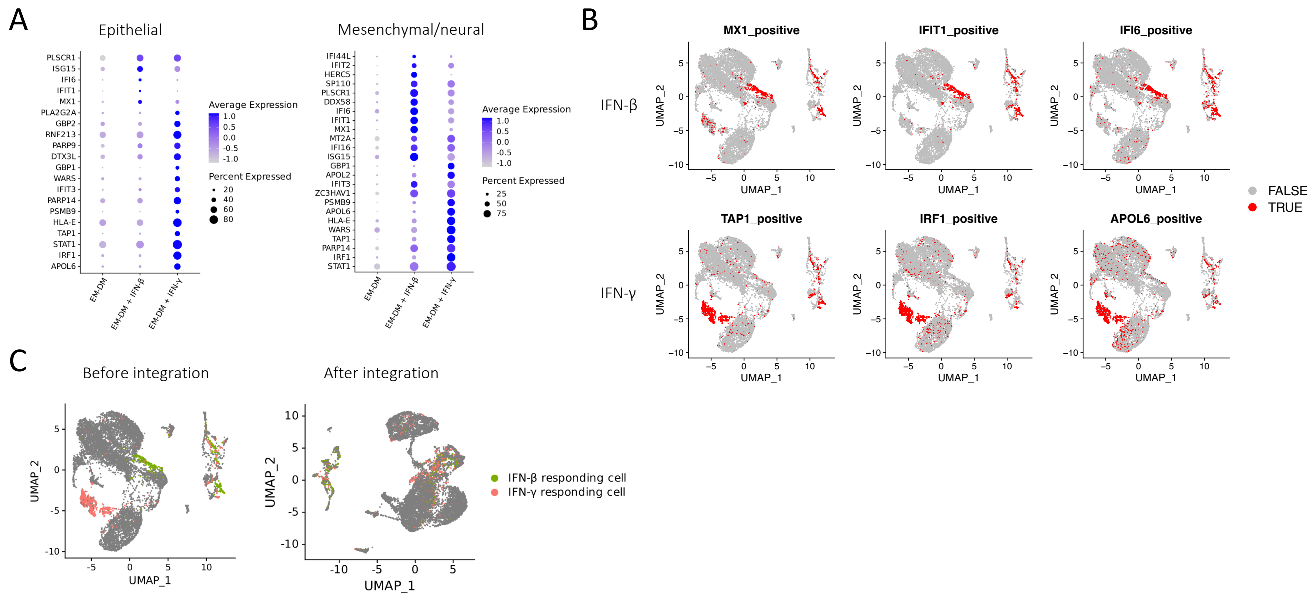


**Supplementary Figure 8. Characterization of the cellular response to IFN-β and IFN-γ in the intestine-on-chip.** (A) Average expression of top differentially expressed genes between the EM-DM and interferon (IFN)-β-stimulated or IFN-γ-stimulated condition in the epithelial and mesenchymal/neural compartment. (B) UMAP projection highlighting the cells positive for the markers used to annotate interferon-responding cells: *MX1, IFIT1* and *IFI6* (IFN-β) and *TAP1, IRF1* and *APOL6* (IFN-γ). (C) UMAP projection highlighting the IFN-β- and IFN-γ-responding cells before and after integration for interferon stimulation.

**Supplementary Table 1. Cell subtype abundance in the different media conditions in the intestine-on-chip.** Cell counts depict the combined counts from three biological replicates.

| Cell type | EM | EM-DM | DM |
|---|---|---|---|
| TA/stem cell | 735 | 340 | 190 |
| Enterocyte progenitor | 313 | 575 | 352 |
| Enterocyte type 1 | 199 | 791 | 1144 |
| Enterocyte type 2 | 144 | 1005 | 507 |
| Paneth-like cell | 306 | 806 | 963 |
| Goblet cell | 1 | 63 | 83 |
| Enteroendocrine cell | 2 | 75 | 66 |
| Mesenchymal-like epithelial precursor | 1 | 116 | 208 |
| Mesenchymal-like epithelial cell type 1 | 385 | 18 | 0 |
| Mesenchymal-like epithelial cell type 2 | 5 | 14 | 20 |
| Dividing mesenchymal/neural cell | 901 | 61 | 6 |
| Myofibroblast | 94 | 186 | 45 |
| Neuron | 161 | 92 | 48 |
| WNT4-positive neural cell | 28 | 28 | 26 |

**Supplementary Table 2. Epithelial cell subtype proportions in the intestine-on-chip compared to the human adult small intestine.**

Percentage of cell types in epithelial compartment, displayed as average values with standard deviation. Data from the human adult small intestine is derived from the Gut Cell Atlas.

| | Intestine-on-chip | | | Human |
|---|---|---|---|---|
| Cell type | EM | EM-DM | DM | Adult small intestine |
| TA/stem cell | 34.9% (SD 2.8) | 9.1% (SD 2.2) | 5.7% (SD 1.6) | 15.3% (SD 13.2) |
| Enterocyte | 27.4% (SD 11.2) | 61% (SD 12.7) | 54.5% (SD 9.6) | 72.4% (SD 14.7) |

| | | | | |
|---|---|---|---|---|
| Paneth(-like) cell | 18.0% (SD 9.5) | 22.7% (SD 12.1) | 29.0% (7.1) | 4.1% (SD 4.9) |
| Goblet cell | 0% | 1.4% (SD 1.5) | 2.2% (SD 1.3) | 1.0% (SD 1.4) |
| Enteroendocrine cell | 0.1% (SD 0.2) | 1.8% (SD 1.3) | 1.7% (SD 1.0) | 0.2% (SD 0.4) |
| Mesenchymal-like epithelial cell | 19.6% (SD 3.2) | 4.0% (SD 0.7) | 7.0% (SD 3.1) | *NA* |
| BEST4+ epithelial cell | 0% | 0% | 0% | 5.7% (SD 5.8) |
| Microfold cell | 0% | 0% | 0% | 0.6% (SD 1.1) |
| Tuft cell | 0% | 0% | 0% | 0.7% (SD 1.0) |

**Supplementary Table 3. Primary antibodies used for immunofluorescent microscopy**

| **Antigen** | **Dilution** | **Host/Isotype** | **Catalogue number** | **Manufacturer** |
|---|---|---|---|---|
| VIM | 1:100 | Rabbit IgG | 5741S | Cell Signaling Technology |
| EPCAM/CD326 | 1:500 | Mouse IgG1 | 2929S | Cell Signaling Technology |
| CHGA | 1:100 | Mouse IgG1 | MA5-13096 | ThermoFisher |
| MUC2 | 1:200 | Mouse IgG1 | ab118964 | Abcam |
| LYZ | 1:200 | Mouse IgG2a | NB100-63062 | Novus Biologicals |
| MKI67 | 1:200 | Rabbit IgG | ab16667 | Abcam |
| RBP2 | 1:100 | Rabbit polyclonal | HPA035866 | Sigma Aldrich |
| ZO-1 | 1:100 | Mouse IgG1 | 33-9100 | ThermoFisher |
| CDX2 | 1:100 | Goat IgG polyclonal | AF3665-SP | R&D systems |

**Supplementary Table 4. Secondary antibodies used for immunofluorescent microscopy**

| **Fluorophore** | **Dilution** | **Species reactivity** | **Catalogue number** | **Manufacturer** |
|---|---|---|---|---|
| Alexa Fluor 488 | 1:250 | Mouse IgG (H+L) | 715-545-150 | Jackson ImmunoResearch |
| Alexa Fluor 647 | 1:250 | Goat IgG (H+L) | A-21447 | ThermoFisher |
| Cy3 | 1:250 | Rabbit IgG (H+L) | 711-165-152 | Jackson ImmunoResearch |

**Supplementary Table 5. Antibodies used for flow cytometry**

| **Antigen** | **Fluorophore** | **Dilution** | **Catalogue number** | **Manufacturer** |
|---|---|---|---|---|
| VIM | Alexa Fluor 594 | 1:50 | 7675S | Cell Signaling Technology |
| EPCAM/CD326 | BUV737 | 1:100 | 748397 | BD Biosciences |
| CHGA | PE | 1:1000 | ab213341 | Abcam |
| MUC2 | PerCP | 1:67 | NBP2-34757PCP | Novus Biologicals |
| LYZ | Alexa Fluor 488 | 1:200 | NB100-63062AF488 | Novus Biologicals |
